# When no answer is better than a wrong answer: a causal perspective on batch effects

**DOI:** 10.1101/2021.09.03.458920

**Authors:** Eric W. Bridgeford, Michael Powell, Gregory Kiar, Stephanie Noble, Jaewon Chung, Sambit Panda, Ross Lawrence, Ting Xu, Michael Milham, Brian Caffo, Joshua T. Vogelstein

**Affiliations:** Johns Hopkins University; Stanford University; Child Mind Institute; Yale University, Northeastern University

## Abstract

Batch effects, undesirable sources of variability across multiple experiments, present significant challenges for scientific and clinical discoveries. Batch effects can (i) produce spurious signals and/or (ii) obscure genuine signals, contributing to the ongoing reproducibility crisis. Because batch effects are typically modeled as classical statistical effects, they often cannot differentiate between sources of variability due to confounding biases, which may lead them to erroneously conclude batch effects are present (or not). We formalize batch effects as causal effects, and introduce algorithms leveraging causal machinery, to address these concerns. Simulations illustrate that when non-causal methods provide the wrong answer, our methods either produce more accurate answers or “no answer”, meaning they assert the data are an inadequate to confidently conclude on the presence of a batch effect. Applying our causal methods to 27 neuroimaging datasets yields qualitatively similar results: in situations where it is unclear whether batch effects are present, non-causal methods confidently identify (or fail to identify) batch effects, whereas our causal methods assert that it is unclear whether there are batch effects or not. In instances where batch effects should be discernable, our techniques produce different results from prior art, each of which produce results more qualitatively similar to not applying any batch effect correction to the data at all. This work therefore provides a causal framework for understanding the potential capabilities and limitations of analysis of multi-site data.

## 1 Introduction

The 21^st^ century has seen the advent of high-throughput techniques for acquiring data on an unprecedented scale. Collection of these datasets often occurs through consortia across different sites, requiring post-hoc data aggregation approaches. These “mega-studies” comprise numerous individual studies, rendering samples sizes substantially larger than any individual study and addressing the small-sample size woes associated with modern big biomedical data (Marek et al., 2022).

Unfortunately, aggregating data across diverse datasets introduces a source of undesirable variability known as a batch effect. Lazar et al. (2013) provides a recent consensus definition of batch effect: “the batch effect represents the systematic technical differences when samples are processed and measured in different batches and which are unrelated to any biological variation.” While these batch effects may not be immediately nefarious, their correlation with upstream biological variables can be problematic (Leek et al., 2010). When biological variables are correlated with batch-related variables, our ability to discern veridical from spurious signals is limited (Akey et al., 2007; Conrads et al., 2004). This problem has “led to serious concerns about the validity of the biological conclusions” (Leek et al., 2010) in data that may be corrupted by these biases; that is, it is unclear whether subsequent detected variability can be attributed to the biological variables, or to the so called “batch effect”. Unfortunately, the qualitative description provided by Lazar et al. (2013) presents limited technical information about how batch effects can be detected or mitigated.

Many approaches model the batch collection or measurement process as a nuisance variable (Johnson et al., 2007; Leek & Peng, 2015; Leek et al., 2010; Pearl, 2010b; Pomponio et al., 2020; Wachinger et al., 2020; Yu et al., 2018). The implicit model justifying these approaches assumes batch effects are associational or conditional, but not causal. Such assumptions are strong, potentially unjustified, and often inappropriate. Two of the most prominent examples of these techniques are `ComBat` and Conditional `ComBat` (`cComBat`) (Johnson et al., 2007). These approaches have demonstrated empirical utility in various genomics and neuroimaging contexts (Pomponio et al., 2020; Zhang et al., 2020); however, it remains unclear when these approaches will be successful, and when they will fail. Specifically, it is unclear when they remove biofidelic variability or fail to remove nuisance variability (Bayer et al., 2022). We still do not know when they produce “wrong answers”, removing desirable signal or failing to remove spurious signal.

In this work, we develop a causal approach to define, detect, estimate, and mitigate batch effects. Our main conceptual advance is modeling batch effects as causal effects rather than associational or conditional effects. Given this structure, we introduce a formal definition of causal batch effects. This formal definition reveals the limitations of (typically inappropriate) assumptions implicit in existing approaches (Rosenbaum & Rubin, 1983, 1985; Stuart, 2010) and provides a theoretical explanation for many limitations of batch harmonization noted in Bayer et al. (2022).

Methodologically, we introduce a simple pre-processing strategy that one can apply to existing techniques for batch effect detection and mitigation. First, to detect and estimate batch effects, we introduce Causal cDcorr (Bridgeford et al., 2023; Wang et al., 2015)—building on modern, non-parametric statistical methods—to estimate and detect the presence of batch effects. Second, to mitigate batch effects, we introduce `Matching cComBat`—an augmentation of the `ComBat` procedure (Johnson et al., 2007)—to remove batch effects while limiting the removal of veridical biological variation. Our proposed techniques introduce the possibility of “no answer” to batch effect correction and detection; that is, the data are insufficient to make a conclusion either way.

We apply these methods to simulations and a large neuroimaging mega-study assembled by the Consortium for Reliability and Reproducibility (CoRR) (Zuo et al., 2014), consisting of more than 1,700 individuals across 27 disparate studies. Our simulations and real data analysis demonstrate that existing strategies can, under many realistic use-cases, experience biases wherein they will confidently produce potentially erroneous conclusions regarding batch effects, where our proposed techniques either produce expected behaviors or avoid inference entirely. This work therefore represents a seminal effort to design methodologies which identify or overcome potential biases in the detection and correction of batch effects from multi-site data.

## 2 Methods

### 2.1 A conceptual illustration of the value of causal modeling of batch effects

Consider a simple example in Figure 1. We observe *n* = 300 measurements in two batches, where one batch (orange) tends to sample younger people, and the other batch (blue) tends to sample older people. The observed outcome of interest is disease state (y-axis), and there is a single potential confounder, which is measured: age (*x-axis*). The solid lines indicate the true distribution governing the relationship between age and disease for both batches, which is unknown in practice. That the data-generating distributions differ indicates that a batch effect is present (red band). Techniques are desired to remove the batch effect given only the data measurements (outcome and covariate pairs, indicated as points). The two rows show two different settings: the top shows a case where the orange batch tends to be larger than the blue before any correction is applied (a positive batch effect), and the bottom shows the reverse (a negative batch effect).

**Figure 1:**
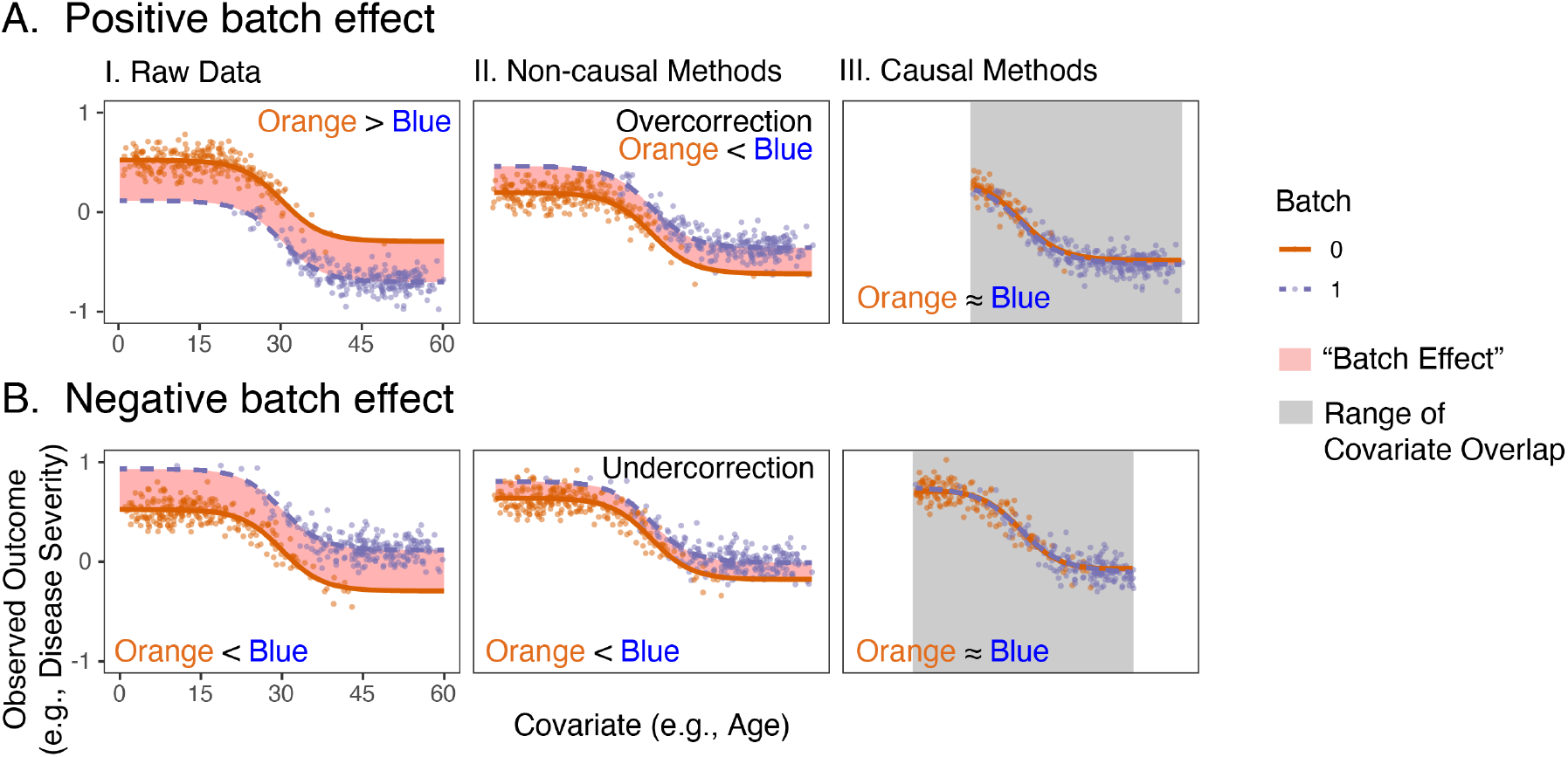
Non-causal batch effect mitigation procedures are subject to both over and under correction and cannot rectify “confounding.” Our causally enriched methods address these issues. **I**. shows the observed data (points), where color indicates batch. The orange line and the blue line indicated the expected outcome per-batch, and the “batch effect” describes the observed difference (red band) between the expected outcomes. Ideally, after batch effect correction, the blue and orange lines should overlap. The orange batch tends to oversample people with younger ages, and the blue batch tends to oversample people with higher ages. **(A)** A scenario where the covariate distributions are moderately confounded and partially overlap, and the orange batch tends to see higher outcomes than the blue batch. **(B)** A scenario where the covariate distributions are moderately confounded and partially overlap, and the orange batch tends to see lower outcomes than the blue batch. **II**. and **III**. illustrate the corrected data after correction, via non-causal and causal methods respectively. If the batch effect is removed, the orange and blue lines should be approximately equal. Non-causal methods attempt to adjust for the batch effect over the entire covariate range, and in so doing, are subject to strong confounding biases. Supposed “batch effect correction” instead introduces spurious artifacts (**(A)**) or fails to mitigate batch effects (**(B)**). Causal methods instead look to a reduced covariate range (gray box), finding points between the two datasets that are “similar,” and are not subject to these biases. Simulation settings are described in Appendix D.2.

Figure 1A illustrates an example with a moderate amount of covariate overlap, meaning that data from the two batches have similar (but not identical) age ranges. This problem is typically conceptualized via the the location/scale (L/S) `cComBat` model of Johnson et al. (2007), which can be generalized as Pomponio et al., 2020:

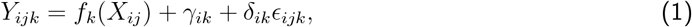

where the measurements *Y*_*ijk*_ for a sample *j* in batch *i* and dimension *k* with covariates *X*_*ij*_ are a linear combination of an overall “true” underlying property *f*_*k*_(*X*_*ij*_) with an additive batch effect *γ*_*ik*_ and a multiplicative batch effect *δ*_*ik*_ for the error *ϵ*_*ijk*_, which is typically assumed to be normally distributed. This model is typically fit via regression, such as the `cComBat` procedure. Non-causal strategies such as `cComBat` learn from each batch, and then *extrapolate* trends across covariates (in this case, age) using the model to infer a relationship between the two batches. The problem is that this approach is strongly sensitive to the specifics of the extrapolation. Because the true data-generating distribution is unknown at the time of analysis, most non-causal approaches perform a linear extrapolation (Chen et al., 2022; Fortin et al., 2018; Johnson et al., 2007; Pomponio et al., 2020), where *f*_*k*_ is assumed to be a linear function. This is typically performed by “removing” the batch effect terms; e.g., the “batch-effect corrected” measurements are:

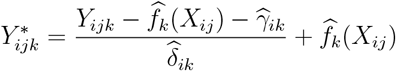

When covariate overlap is imperfect, however, misspecification of *f*_*k*_ can be disastrous (Figure 1A.II). While before correction the true data-generating distribution for the orange batch is higher than the blue batch, after correction the blue batch is higher the orange batch (the batch effect was *over-corrected*; i.e., too much correction was applied). Reversing the relationship between the blue and orange batches in Figure 1B, non-causal strategies are still unable to properly remove the batch effect and may still over-or under-correct for batch effects. As a result, “batch-effect-corrected data” from non-causal strategies may not actually be corrected, and in many situations may be more different after “correction” than before. In other words, even though a fundamental desiderata of batch effect correction is to decorrelate the relationship between the batch and upstream covariates of interest (Leek et al., 2010), the presence of such a correlation also hampers our ability to remove the batch effect.

On the other hand in Figure 1A.III or Figure 1B.III, causal techniques focus instead on deriving conclusions within a range of covariate overlap in the data, where confounding is better controlled. The true data-generating distributions (and the points themselves) almost perfectly align after batch effect correction across the range of overlapping covariates, indicating that the batch effect has been removed, despite no foreknowledge of the true data-generating model.

In this manuscript, we posit that we desire that methods report confounding when it is present, something only causal methods do. Motivated by this perspective, we augment traditional approaches with causal machinery for batch effect detection and correction. Theory, simulations, and real data analysis demonstrate that traditional strategies to detect or correct batch effects from mega-studies lack the ability to identify confounding, and therefore, often add or remove batch effects inappropriately when covariate overlap is low. Therefore, such approaches cannot be trusted without further analysis. These issues highlight the primary challenges of performing valid statistical inference while pooling data across studies. This work therefore contributes to the ongoing effort to improve the validity and reliability of inferences in past and future mega-studies.

### 2.2 Models motivating different batch effect techniques

Here we build up the implicit modeling assumption underlying various approaches to mitigating batch effects. Our working example is that the exposure is batch, and the outcome is connectome. We want to know whether the differences between the batches are due to variability in the participants themselves who are measured, or veridical differences in outcomes across the two batches. Figure 2 shows four different models and provides details of which kinds of variables are relevant in neuroimaging studies, though the concepts apply much more generally. Directionality of arrows (e.g., *T* → *Y*) indicates that the variable *T* can influence the variable *Y*. Therefore, our goal is to identify a regime in which a strategy can provide evidence of a batch effect; that is, the degree to which the exposure influences the outcome (i.e., a *causal* effect).

**Figure 2:**
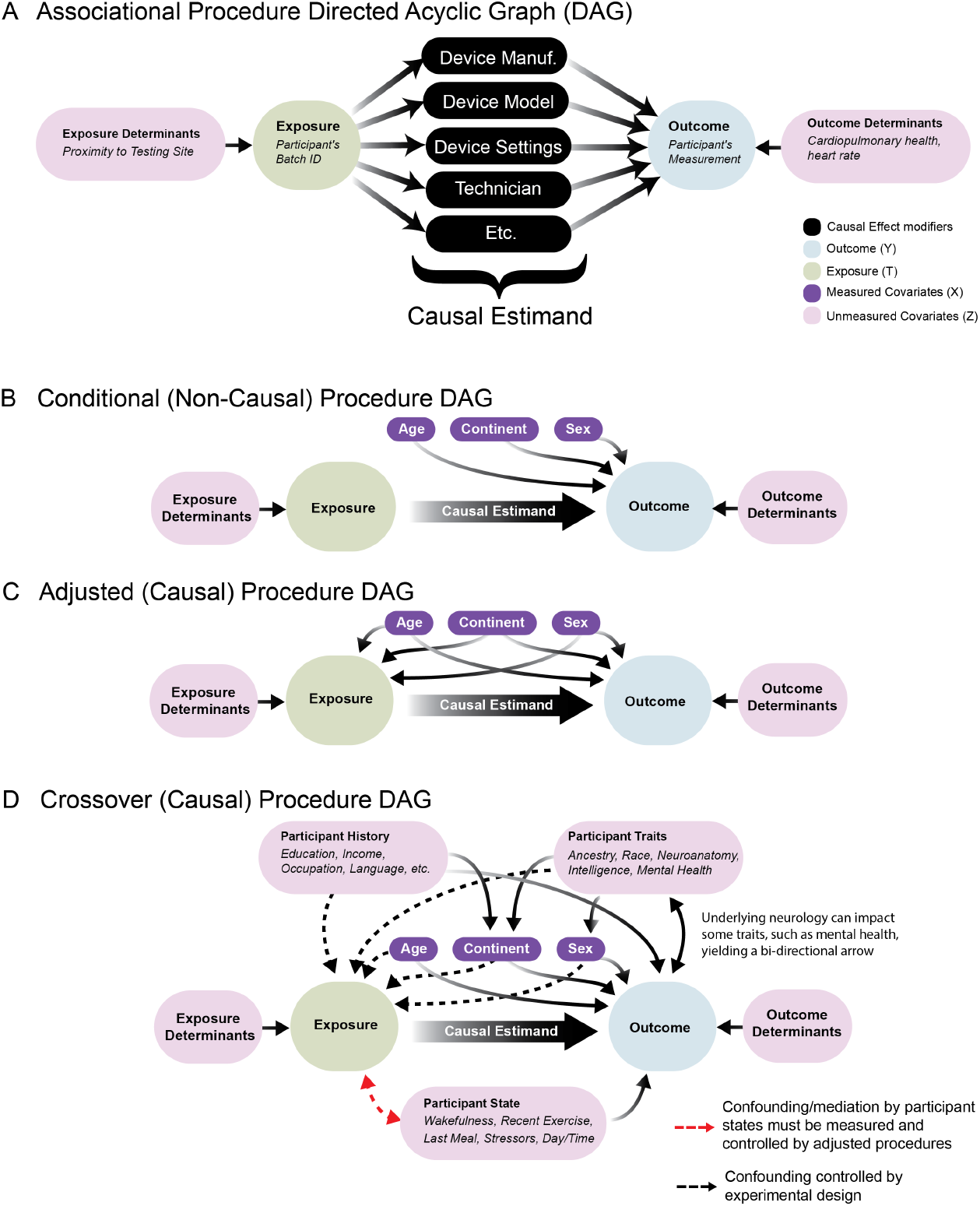
Causal Graph of Study Covariates. Causal Directed Graphs illustrating the underlying assumptions under which various procedures to detect or correct for batch effects are reasonable. Boxes represent variables of interest, and arrows indicate “cause and effect” relationships between variables. The causal estimand is the impact of the exposure (the batch) on the outcome (the participant’s measurement) and is a cumulative effect of effect modifiers (black variables) both known and unknown that yield batch-specific differences. The relationship between the exposure and the outcome are *confounded* if there are open backdoor paths (Pearl, 2009b). **(A)** Associational procedures and **(B)** conditional procedures are reasonable when there is no confounding. **(C)** Adjusted causal procedures are reasonable when backdoor paths can be blocked by measured covariates (Pearl, 1995, 2009b). **(D)** Crossover procedures are reasonable under many forms of potential confounding, both measured or unmeasured, so long as participant states are non-confounding. If participant states are confounding, the confounding states must be measured (red arrow). Note that some participant traits (e.g., intelligence or mental health) may be caused by the connectome, and introduce the potential for bi-directional arrows in the causal graph with the measurement, visited in the Discussion Section 4.

We begin with the simplest approach, an “Associational” model (Figure 2A). Using an associational model enables us to obtain valid answers to: “are differences in outcome associated with differences in exposure?” The answer to this question is often important, for example, in biomarker studies, but may be insufficient for determining causality. Dcorr(Székely et al., 2007) and naive `ComBat`(Johnson et al., 2007) (e.g., the model in Equation (1), but omitting covariate modeling via *f*_*k*_(*X*_*ij*_)) are methods designed to test and correct for associational effects. For example, if the batch and the outcome are dependent upon any covariates, which they often are in real studies, then an estimated associational effect is not a valid estimate of a batch effect (Figure 2**(A)**). For instance, if the two batches differ because they have sampled individuals in different age groups, and the outcome is a function of age, then using this approach will lead to falsely concluding causal batch effects. On the other hand, if a batch effect is truly present, but somehow another covariate cancels out this variability, then this approach could falsely conclude there does not exist a batch effect when it is present.

The “Conditional” model (Figure 2B) enables us to seek valid answers to a more nuanced question: “are differences in outcome associated with differences in exposure, after conditioning on measured covariates (Figure 2B)?” If there are differences in outcome after conditioning, this can suggest that the differences are due to different exposures. cDcorr(Wang et al., 2015) and `cComBat`(Johnson et al., 2007) (e.g., the model in Equation (1)) are methods designed to test and correct for conditional effects. However, this model is subject to errors in the presence of confounding. The covariates might also impact the exposure, and if so, also impact our ability to identify a causal estimand, but without our knowledge. Imagine, for instance, that the two batches have covariates that partially overlap; for example, one sampled adolescents and adults, but not geriatrics, and the other sampled adults and geriatrics, but not adolescents. Conditional approaches make implicit assumptions about how to interpolate outcomes across covariate distributions potentially unobserved in a given batch. When those assumptions do not reflect the real data, the results can be erroneous. In this light, the identified effect may be a veridical batch effect, or an artificial covariate effect, thereby leading to either false positives or false negatives.

The “Adjusted” model (Figure 2C) enables us to mitigate this concern. Here, we seek valid answers to the following question: “conditioned on any impact of measured covariates on either exposure or outcome, do we see any residual differences in outcomes associated with differences in exposure?” This is effectively the model in most observational studies, which are a critical component of causal inference (Stuart, 2010). Our proposed Causal cDcorr and `Matching cComBat` (described below) are methods designed to test and correct for data following this model (Bridgeford et al., 2023). For example, to address the above concern, an adjusted model might discard all the adolescents and geriatrics, and simply focus on modeling the adults, for which we have data in both batches. While this reduces the sample size, it avoids the pitfalls of confidently concluding the presence or absence of a batch effect, even when it is not clear. Further, conclusions readily generalize to samples similar to the reduced samples, so many of the excluded samples may still be able to be batch effect corrected and used in subsequent inference. However, it also suffers from a potential confounding issue. While adjustment can be used to address confounding with measured covariates, it cannot be used to address confounding with *unobserved* covariates. Specifically, if there exist unobserved covariates that do not correlate with the observed ones and yet impact the outcomes, exposures, or both, then adjusted methods may lead to spurious results.

The “Cross-over” model (Figure 2D) addresses this concern. In a cross-over investigation, each participant is measured across *all* exposure groups. When properly performed, cross-over models enable us to seek valid answers to questions such as: “conditioned on almost all potential confounding, are differences in outcome associated with differences in exposure?” When the experimental design is a cross-over study and states are unchanging or randomized, simple `ComBat` approaches can be adequate, though the authors are not aware of any extensions of distance correlation or other non-parametric tests to these paired settings for high-dimensional data. If participant states change, then confounding remains unless the order participants are measured in each batch is randomized or participant states are measured and suitably adjusted for, as-per the above adjusted model.

The importance of these models can be formalized more rigorously through the statistical notions of causal consistency, positivity, and ignorability. Reasoning that data may satisfy these assumptions forms the cornerstone of causal inference from observational studies. In the context of an investigation with batch effects, *causal consistency* asserts that each individual has a set of potential measurements corresponding to each possible batch, and the observed measurement for an individual is equivalent to their potential outcome for the batch they were actually measured in (Frangakis et al., 2007). This criterion can be intuitively be reasoned for batch effects investigations, in that individuals could conceptually be measured across all batches (but we only observe their potential measurement for the batch in which they were measured). *Positivity*, equivalent to *covariate overlap*, states that for every combination of covariates, there must be a non-zero probability of being measured in any of the included batches (Rosenbaum & Rubin, 1983, 1985). Adjusted techniques attempt to enforce positivity or covariate overlap on the data, by reducing the scope of inference to individuals who could have been measured across all batches. *Ignorability* asserts that given the observed covariates, the batch assignment and the potential outcomes are independent. Stated another way, conditional on the observed covariates, the batch assignment process is effectively random, and there are no unmeasured factors that influence *both* batch assignment and the potential outcomes (Rosenbaum & Rubin, 1983). While this may seem like an insurmountable hurdle, measured covariates are often sufficient conditioning sets so long as the measured covariates are suitably correlated with unmeasured variables (Pearl, 2009a). Appendix A provides rigorous definitions and mathematical models contextualizing the above definitions.

### 2.3 Detecting and mitigating batch effects

The causal graphs in Figure 2 of batch effects makes immediately clear their undesirability. If we want to learn about the impact of any upstream variable on the measurement, *batch effects may be a mediator of the relationship between that variable and the measurement*. If we want to learn about the impact of the measurement on any traits of a person (such as in a brain-behavioral investigation), *batch effects may introduce cycles in the causal graph, due to the the inability to faithfully represent underlying neurological properties free from such measurement errors*. Both of these characteristics are immediately problematic for subsequently deriving causal conclusions, as failure to account fort the batch effect may limit the identifiability of other potential estimands we may wish to subsequently investigate (Pearl, 2009b, 2010a, 2014). Removal of the batch effect on the data derivative represents a strategy to control for these problematic characteristics (via deletion of the arrow on the causal graph from batch to measurement), and motivates why studying, understanding, and deciphering strategies to detect or remove batch effects are desirable for subsequent inference tasks. Colloquially, this causal presentation delineates that batch effects are not just undesirable artifacts that could potentially yield spurious correlations (Leek et al., 2010), but asserts that a failure to account for them *may prohibit principled subsequent inference*.

At present, literature approaches for detecting and mitigating batch effects tend to (implicitly or explicitly) model the effects as associational or conditional, which often fail to adequately account for confounding biases. To this end, we propose a simple technique to augment classical strategies (Figure 3, see Appendix C for methodological details):

**Figure 3:**
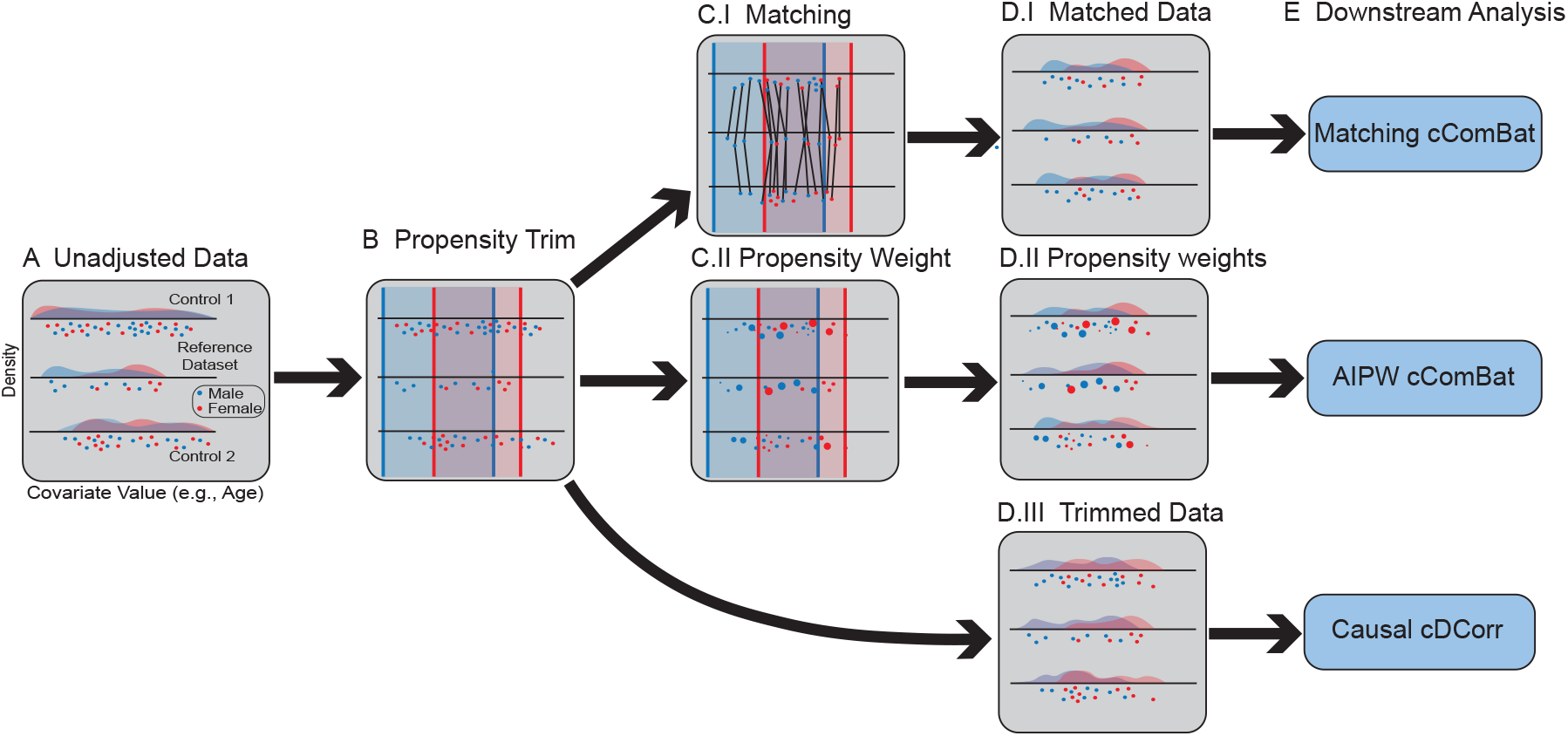
The demographic balancing procedure serves to demographically align poorly balanced datasets using causal approaches. **(A)** The unadjusted datasets are imbalanced in covariate distributions. The *reference dataset* is indicated. **(B)** Propensity trimming (shaded boxes) provide general alignment of demographics, such that no datasets will include demographics unrepresented in other datasets. **(C.I)** samples from other datasets are matched to samples from the reference dataset, and **(D.I)** samples without matches are discarded from subsequent analysis. The adjusted data after matching have nearly identical covariate distributions. **(C.II)** samples are weighted according to their inverse propensities, such that samples which look non-representative of other datasets are down-weighted. **(D.II)** This can serve to also yield somewhat similar covariate distributions after re-weighting. **(D.III)** the trimmed data generally feature overlapping covariate distributions, but may not be identical across datasets. **(E)** Downstream analysis for batch effect correction or detection applied to the adjusted data (and potentially propensity weights) via `Matching cComBat`, `AIPW cComBat`, or Causal cDcorr.

1. Use classical causal procedures to re-weight the measurements from the batches so that measured covariate distributions across all batches are overlapping or balanced. For batch effect detection, we use vector matching (Lopez & Gutman, 2014), a form of propensity trimming which performs a multinomial regression of batch onto the covariates and then trims individuals with uncharacteristically low or high estimated probabilities (*propensities*) for any of the batches. For batch effect correction, `Matching cComBat` uses vector matching followed by a nearest neighbor matching (without replacement) of a “reference study” to the other studies in a mega-study. This matching may be performed many-to-one or one-to-many depending on the relative sizes of the reference study and the other studies. We propose the use of exact matching on categorical or binary covariates when possible, and Mahalanobis distance matching for continuous covariates. Our experiments and real data use-cases leverage a 0.1 distance caliper, which upper-bounds the Mahalanobis distance between a given matched pair of individuals.
2. Apply traditional procedures for detection or correction of batch effects post re-weighting. We use cDcorr for batch effect detection (Bridgeford et al., 2023; Wang et al., 2015), and `cComBat` for batch effect correction (Johnson et al., 2007; Leek & Peng, 2015; Leek et al., 2010; Pearl, 2010b; Pomponio et al., 2020; Wachinger et al., 2020; Yu et al., 2018).
3. Optionally, apply estimated batch effect corrections to excluded samples with covariates similar to the re-weighted samples or samples demographically dissimilar to the re-weighted samples. The former requires few additional assumptions, and may increase sample sizes for subsequent inference. Applying learned batch effects to demographically dissimilar individuals requires stringent assumptions regarding whether extrapolations to individuals demographically dissimilar from the re-weighted samples is appropriate.

Our proposed augmentations via classical causal procedures could be equivalently exchanged for other re-weighting procedures, such as inverse-probability weighting (IPW) or augmented inverse-probability weighting (AIPW). Our simulations explore an additional technique, `AIPW cComBat`, which uses augmented inverse-probability weighting to produce batch effect estimates. Through AIPW, we estimate propensity weights from the observed data, and incorporate these propensity weights into outcome model regression (J. Robins, 1986). While not the focus of this article, we explore the utility of these methods in simulation settings.

## 3 Results

### 3.1 Causal machinery helps mitigate limitations of traditional batch effect methods

Here we quantify the performance of causal and non-causal methods by generalizing the simulations from Figure 1. 1000 samples are generated from one of two batches. Figure 4A shows multiple simulation settings, including linear (left), nonlinear (center), and non-monotonic (right). Solid lines indicate the expected outcome at a given covariate value for a particular batch. Appendix D provides details for the empirical settings of the simulations. The standard measure of effect size in the causal literature is called the ‘Average Treatment Effect’ (ATE) (Rosenbaum & Rubin, 1983, 1985). The red bands in Figure 4A show the treatment effect for all possible values of the covariate, so the ATE is just the average width of that bar, which happens to be −1 here. In this case, we consider the Average Absolute Treatment Effect, which we denote by ‘AATE’ for brevity, which is the average absolute width of the bar, which happens to be 1 here. Since the treatment here is batch, the goal is to remove the treatment effect, which requires correctly estimating the treatment effect and then removing it. When AATE = 0 (no batch effect), the covariate/outcome relationship is equal across the two batches, and non-zero otherwise.

**Figure 4:**
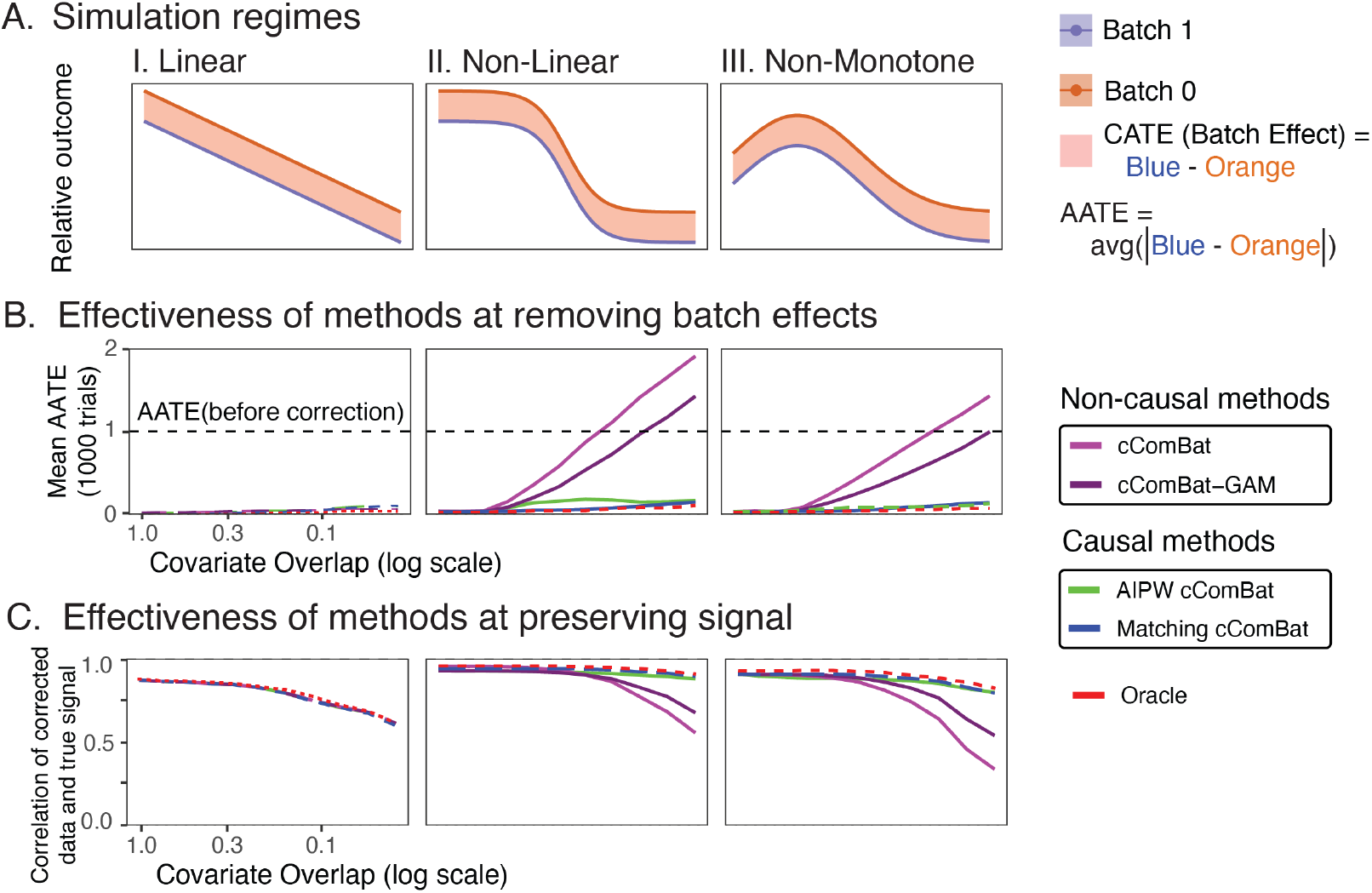
Simulation regimes illustrate that non-causal procedures are subject to strong biases without covariate matching. **(A)** illustrates the relationship between the relative expected outcome and the covariate value, for each batch (color), across **(I.)** linear, **(II.)** non-linear, and **(III.)** non-monotone regimes. The conditional average treatment effect (red box) highlights the batch effect for each covariate value. The average treatment effect (ATE) is the average width of this box, and the average absolute treatment effect (AATE) is the average absolute width of this box. In these simulations, the AATE before treatment is 1. **(B)** The effectiveness of the techniques at removing the batch effect. Techniques with high performance will have a mean AATE after correction at or near 0 (the batch effect was eliminated). **(C)** illustrates the effectiveness of different batch effect correction techniques for preserving the underlying true signal. Techniques with high performance will have higher correlations with the underlying true signal.

We consider four different algorithms for this purpose: `cComBat`, `cComBatGAM`, `Matching cComBat`, and `AIPW cComBat`. These approaches are compared to the `oracle`, a batch effect correction technique which has prior knowledge of the true data generating model. We show the estimates of the mean AATE after correction over 1000 trials for each method and each simulation setting as we vary the amount of covariate overlap (top row, Figure 4B). Ideally, after batch effect correction is performed, the covariate/outcome relationship will be equal across the batches, and the AATE will be approximately zero. When the relationship between covariate and outcome is linear, all the methods work regardless of the amount of overlap (Figure 4C.I). However, when the relationship is non-linear, the non-causal methods mis-estimate the batch effect unless there is a nearly perfect overlap, and the resulting data are qualitatively dissimilar across batches after correction, indicated by the mean AATE being far from 0 (Figure 4B.II). In cases of extreme non-overlap, note that the data are similarly dissimilar after batch effect correction to before any correction were applied (AATE near or above the dotted black line). In contrast, `Matching cComBat` and `AIPW cComBat` correctly estimate and remove the batch effect, and demonstrate near optimal performance of the `oracle`. The results are qualitatively similar when the relationship is non-monotonic (Figure 4B.III).

In addition to removing batch effects, it is critical that a batch effect correction technique preserves underlying signal in the data. We evaluate how well the corrected data reflects the true underlying relationship (linear, non-linear, or non-monotonic) between the covariate and the outcome. We compare the corrected data to the true relationship using Pearson’s correlation Pearson, 1896, restricting to the matched samples so that the correlations are all computed with respect to the same set of sample in Figure 4(C). Low correlations indicate that the data poorly reflect the true relationship, suggesting that regardless of whether or not there is a batch effect, the underlying signal has been perturbed. Again, regardless of the *a priori* presence of batch effects, causal methods tend to outperform non-causal methods for preserving the underlying signal in the data, and show performance near that of the `oracle`, particularly as covariate overlap declines.

The results are qualitatively and quantitatively similar when there is no batch effect *a priori* (the focus of Appendix D.5), indicating that non-causal methods can also introduce artifacts to the data when none are present. On the other hand, causal methods show greater robustness to these situations too, with performance closely mirroring the `oracle`. Taken together, these results suggest the utility of causal methods for identifying and correcting for batch effects from data, as they have near-optimal performance in our simulation regimes for identifying and removing batch effects when batch effects are present, and avoiding the introduction of artifacts to the data when no batch effects are present *a priori*.

### 3.2 The CoRR Studies have disparate demographic characteristics

The motivation for our work is the neuroimaging mega-study produced by the Consortium for Reliability and Reproducibility (Zuo et al., 2014), a collection of over 3,500 functional neuroimaging measurements from over 1,700 individuals spanning 27 separate datasets. A full description of data pre-processing for the neuroimaging and covariate information is provided in Appendix E. Figure 5A explores the demographic characteristics for the individuals in the CoRR mega-study. Many of the studies have a narrow age range, and several studies only include females. Because sex (Ingalhalikar et al., 2014; Satterthwaite et al., 2015; Weis et al., 2020), age (Hampson et al., 2012; Sala-Llonch et al., 2015; Varangis et al., 2019), and continent (as a surrogate for race and culture) (Ge et al., 2023; Misiura et al., 2020) are variables that have been associated with brain connectivity, they serve as measured demographic covariates used in our investigation.

**Figure 5:**
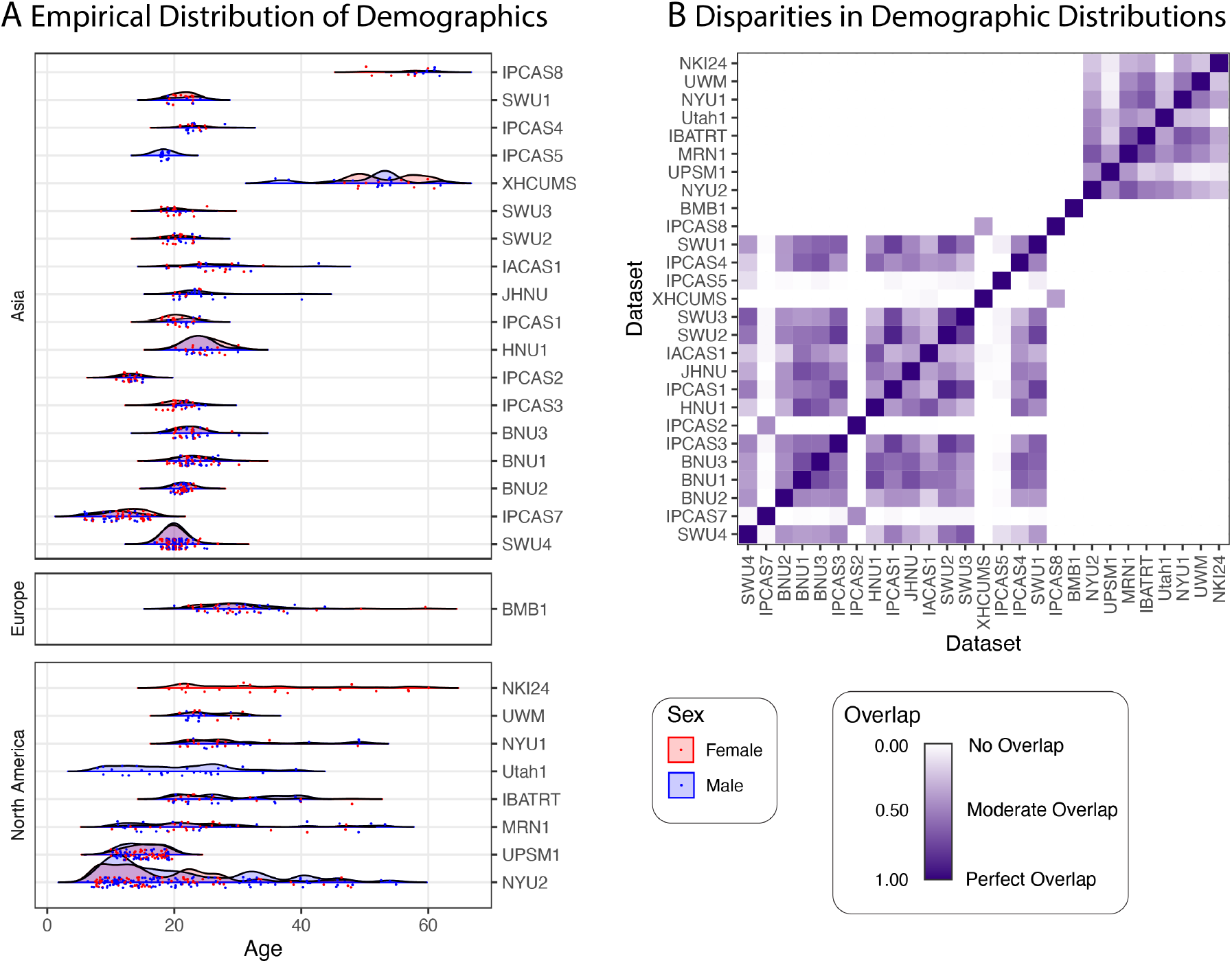
Demographic data for the 27 studies from the CoRR Mega-Study. **(A)** Each point represents the age of a participant corresponding to a single measurement. Rows are studies, boxes are continents, and color indicates sex. *n* = 3,597 samples are shown which featured age, sex, and continent information, and were successfully processed to connectomes. **(B)** Even with only 3 observed covariates (sex, age, and continent of measurement), the CoRR studies often show extremely limited covariate overlap (Pastore & Calcagnì, 2019). This makes inference regarding batch effects difficult.

Figure 5B illustrates the level of demographic overlap in the CoRR mega-study, using a distribution-free overlapping index (Pastore & Calcagnì, 2019) (see Appendix E.2 for details). The CoRR mega-study includes many pairs of datasets with varying degrees of overlap, from high (near 1) to low (near 0). Further, many of the datasets do not overlap at all (overlap of 0), making inference about batch effects impossible without making strong assumptions. This is particularly troublesome, as the covariate records for the CoRR mega-study common to all sites include only three covariates: age, sex, and continent of measurement. Additional measured covariates can only reduce the estimated overlap between the pairs of datasets, so having poor overlap on such a sparse set of covariates indicates that the actual demographic overlap is likely even lower.

### 3.3 Detecting Batch Effects in the CoRR mega-study

Figure 6A focuses on discerning the viability of different types of effects one could use to test for batch effects from Section 2.2. For each pair of datasets in the CoRR study, we test whether (or not) a discernable effect is present, while controlling for age, sex, and continent differences between the individuals within each dataset. Intuitively, we would like to believe that if such a test rejects the null hypothesis in favor of the alternative, the data supports that a batch effect is present. When we account for demographic covariates using conditional (non-causal) approaches (orange squares, top left), differences between the datasets are detected 26.9% of the time (checkboxes, cDcorr, BH correction, *α* = 0.05). Conditional procedures may be “confounded” in extreme cases; e.g., the batch effect conditional on age and sex is confounded between NKI24 and IPCAS5 because one dataset is entirely female and the other is entirely male, and the age distributions do not overlap (Figure 5(B)). This results in cDcorr being unable to compare the conditional distributions (across batch), since the conditional distributions are non-comparable across the batches in the data. Further, as explained in Section 2.2, the viability of these approaches as tests for batch effects is the assumption that the measured covariates are overlapping and non-confounding, which is overwhelmingly false for many of these comparisons, as shown in Figure 5B. Similar tests are performed using adjusted (causal) approaches in 6A (blue squares, bottom right). Many pairs of studies could not have an adjusted (causal) effect estimated due to poor demographic alignment (242 of 351 pairs of studies), where here “poor demographic alignment” corresponds to fewer than 30 samples retained after matching across the two sites. Notably, adjusted (causal) procedures can be used to estimate effects between all pairs of the American Clique (a group of separate studies consisting of similar demographics collected in North America, consisting of NYU1, NYU2, IBATRT, UWM, MRN1). After adjustment for the sample characteristics of the demographics associated with individual studies, a majority of adjusted (causal) effects (66.1%) remain significant (Causal cDcorr, BH correction, *α* = 0.05).

**Figure 6:**
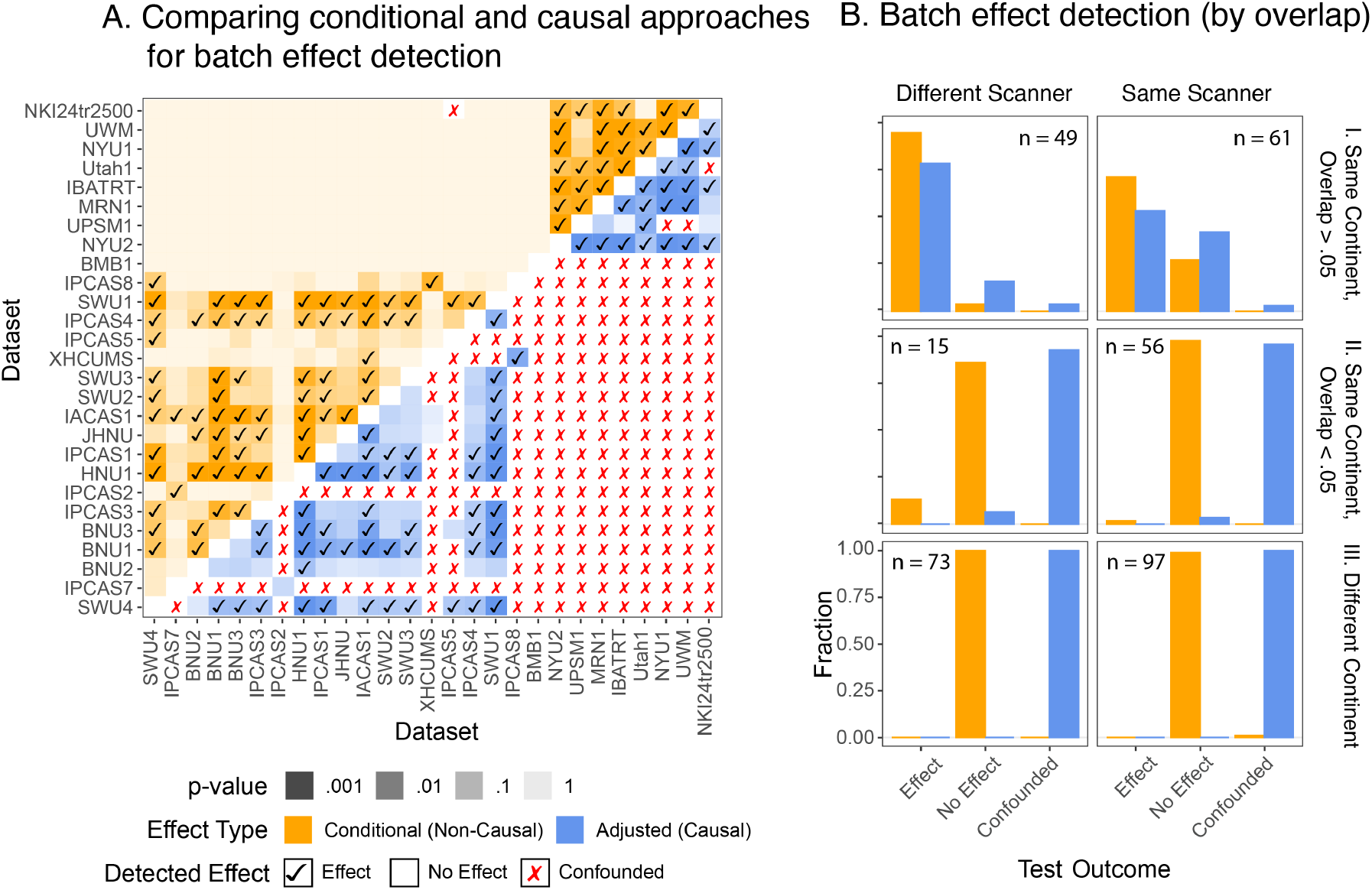
Comparison of types of effects between datasets from the CoRR study. **(A)** Heatmap of different types of effects (conditional and adjusted causal procedures) that can be used to detect differences between each pair of datasets in the CoRR study. Whereas most (25.8%) non-confounded conditional effects are not significant, most (66.1%) non-confounded adjusted effects are significant. **(B).I** delineation of how one possible source of batch effects, scanner model, impacts significance rates of batch effects. Almost all pairs of studies conducted on different scanners with high covariate overlap (*>* .05) have discernable batch effects. The rate of detected effects is lower when the scanner model is the same. **(B).II** and **(B).III** When the level of estimated covariate overlap is lower (*<* .05) or zero (different continent), conditional effects never detect a difference across datasets. On the other hand, adjusted causal procedures instead report that the data is too confounded for subsequent inference and avoid running entirely.

Note that confounding (red “X”s) in Figure 6A for causal effects extremely closely align with datasets with low covariate overlap in Figure 5B. A possible source of a batch effect could be different types of scanners were used to collect the data (Chen et al., 2022; Johnson et al., 2007; Pomponio et al., 2020), which is represented in our presentation as a causal effect modifier in Figure 2A. To this end, we aggregate the fraction of comparisons which report the indicated test outcome for different levels of covariate overlap (rows) when the scanner model is the same (right column) or different (left column). When the scanners are different and the covariates overlap, both causal and non-causal methods reliably detect batch effects a majority of the time, and they do so at a higher rate than when the scanner is the same (Figure 6B.I). When the covariate overlap is low (Figure 6B.II), or when the continent of measurement is different entirely (and the overlap is zero, (Figure 6B.III), conditional (non-causal) procedures almost always fail to reject. On the other hand, adjusted (causal) procedures overwhelmingly report that a given pair of studies are demographically confounded, and are too different for any effect to be investigated.

Appendix E.3 computes numerous within-individual topological properties of functional connectomes, illustrating that `ComBat`-derived approaches do not tend to disrupt within-individual signal of the connectomes, which leaves open the possibility that while within-individual signal may be preserved, cross-individual variability may be altered by the techniques, and motivates our final analysis.

### 3.4 Traditional Approaches Produce Disparate Inference from Techniques Leveraging Causality

We investigate the strength of the sex effect, conditional on age, in the connectomes before and after batch effect correction (Figure 7). Connectomes are filtered to the American Clique, a selection of datasets with overlapping demographics from Figure 5(A), and then either have no batch effect correction, `ComBat`, or `cComBat` (the non-causal approaches) applied to the resulting derivatives, and are finally filtered to a *matched subset* of individuals (Figure 7A). Similarly, we apply `Matching cComBat` to the American Clique, which conceptually applies batch effect correction to the matched subset itself. In this sense, while the manner in which the data were post-processed for batch effects differs, the actual individuals included and the techniques used for the subsequent analysis are identical. We test for a significant sex effect, conditional on age, using the generalised covariance measure. The generalised covariance measure (Shah & Peters, 2018) is a conditional independence test, which performs nonlinear regressions of the outcomes (connectivity) on the conditioning variable (age) and then tests for a vanishing covariance between the resulting residuals across sex. These tests can be adapted for two-sample testing regimes, such as for testing for differences across sex (Panda et al., 2025). The edges which show a significant sex effect conditional on age (generalized covariance measure, BH correction, *α* = .05) are colored from smallest test statistic to largest test statistic (rank = 1, dark purple), and edges without a significant conditional sex effect are colored white (Figure 7B).

**Figure 7:**
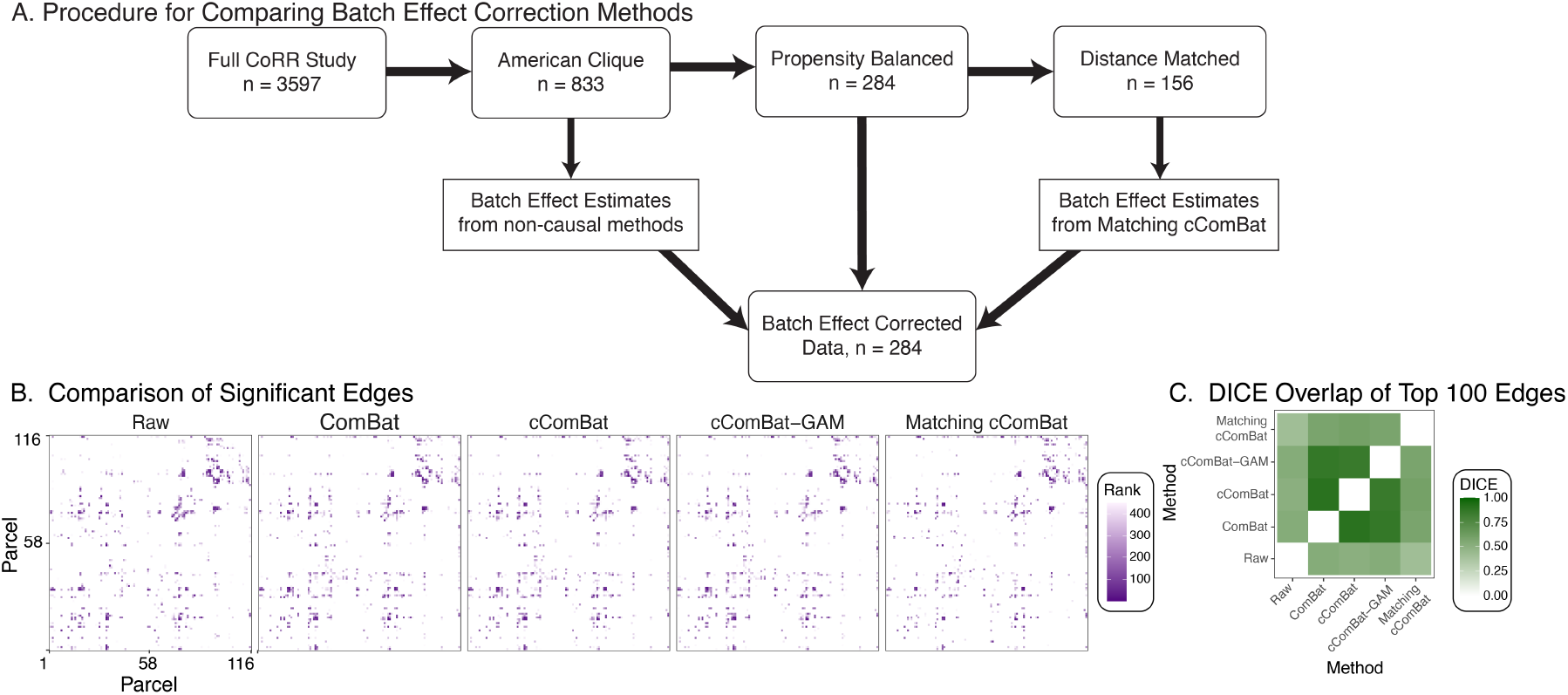
Significant Edges Before and After Batch Effect Removal. **(A)** The presence of a sex effect (conditional on individual age) is investigated for each edge in the connectome. Significant edges are shown in rank order from largest (rank = 1) to smallest sex effect (*α* = .05, Benjamini Hochberg (Benjamini & Hochberg, 1995) correction). Analysis is performed on the American Clique, and then either pre-hoc filtering (e.g., before any batch effect correction is applied, in the case of `Matching cComBat`) or post-hoc filtering (e.g., after batch effect correction is applied, in the case of Raw, `ComBat`, `cComBat`) to a matching subset of individuals. Subsequent inference appears qualitatively dissimilar, even though the subjects used for inference are the same. **(B)** the DICE overlap of the top *n* = 100 edges, by effect size, between all pairs in **(A)**. `cComBat`, `ComBat`, and no batch effect correction provide similar subsequent inference, whereas `Matching cComBat` provides disparate subsequent inference.

While these plots appear superficially similar, we compare the DICE overlap (Dice, 1945; Sørensen, 1948) of the top 100 edges (by effect size) across all approaches. A high DICE overlap indicates that the subsets of edges are similar. `ComBat`, `cComBat`, and `cComBatGAM` tend to produce similar inferences. In contrast, they all have lower overlap with `Matching cComBat` (Figure 7C), and all produce more similar subsequent inference to the raw connectomes (no batch effect correction) than `Matching cComBat`. This suggests that causal and non-causal strategies for batch effect correction can yield dissimilar subsequent inference, even though the selection of individuals being subsequently analyzed is the same.

## 4 Discussion

Succinctly, this manuscript is driven by two questions: (1) should data collected from mega-studies be combined or analyzed separately, and (2) when it should be combined, the optimal strategies for doing so. Our primary conceptual contribution is establishing that batch effects can be formalized as causal effects. This formulation explicitly delineates the desirability of detecting and controlling for batch effects, as failure to do so limits the identifiability of other potential estimands of interest one may wish to study. Given this realization, we propose a framework for quantifying and removing batch effects in high-dimensional datasets featuring non-Euclidean measurements. We propose augmenting existing strategies for proper batch effect correction and removal by prepending causal balancing procedures.

We explore the advantages of these procedures by demonstrating that causal approaches (such as `Matching cComBat` and Causal cDcorr) yield empirically valid approaches to handling batch effects, or report that no inference can be made due to insufficient covariate overlap. This represents a principled approach over alternative strategies, which we show may introduce spurious biases and dramatically under-or over-estimate batch effects. We illustrate how, ignorant of potential confounding, otherwise principled strategies may misrepresent batch effects. This demonstrates that causal adjustment procedures can augment hypothesis testing by indicating when the data may be unsuitable to provide answers to a particular question due to irreconcilable confounding, rather than permitting false positives or false negatives.

Our neuroimaging use-case shows the utility of the methods developed herein for estimating and removing batch-related effects from large biological datasets. We demonstrate that many non-detected effects in the CoRR study may yield false conclusions due to excessive confounding between datasets. When demographic overlap was high and the scanner differed, batch effects were nearly always detectable using conditional (non-causal) or adjusted (causal) procedures. Conversely, with lower demographic overlap, effects were entirely undetectable by conditional procedures. Adjusted (causal) procedures explicitly delineate when the datasets are overly confounded by demographic covariates. We believe this to be a more desirable conclusion, as it highlights that in many cases, even with infinite data following similar covariate distributions, determining the presence or absence of a batch effect would be impossible due to high demographic confounding.

We conclude that batch effect correction represents a theoretically appropriate step supported by simulations for mega-studies with sufficient covariate overlap. When such overlap exists, causal methods demonstrate substantial performance improvements over alternative methods for eliminating undesirable spurious variability without distorting underlying veridical signal. Our methods identify different downstream demographic effects compared to prior approaches, even though the same individuals were analyzed, whereas prior approaches produce conclusions that are qualitatively more similar to the unharmonized data than causal approaches. These approaches are established in the causal literature to permit numerous false positives and false negatives (Ding & Miratrix, 2015; Forstmeier et al., 2017). Together, our work rigorously defines and studies measurement bias and strategies to mitigate demographic confounding in complex, non-Euclidean data.

## Limitations

Researchers may perceive a tradeoff: when datasets lack high demographic overlap, model limitations can impart substantial bias on the investigation’s conclusions. Conversely, enforcing demographic overlap through propensity trimming or matching may seem to yield fewer samples or narrow the scope of conclusions. In practice, with biological data lacking ground truths, we cannot evaluate the introduction of imparted residual bias or determine whether downstream conclusions stem from these biases.

We do not claim that our specific choices for re-weighting measurements are ‘optimal’. Rather, we chose simple, principled approaches for illustrative purposes. Our proposed methods can be conceptualized as “bias-corrected” matching approaches, where we combine matching approaches as a non-parametric data pre-processing step with subsequent regression adjustments (Abadie & Imbens, 2011; Rubin & Thomas, 2000). These methods have been shown to reduce model dependence in parametric settings, such as those leveraged by `cComBat` or `cComBatGAM` (Ho et al., 2007). Our simulations elucidate that even these simple balancing approaches sufficiently highlight substantial shortcomings in existing approaches for detecting or correcting batch effects, and that causal machinery may mitigate these shortcomings.

In our real data analysis, we used only three covariates: age, sex, and continent of measurement (as a surrogate for race) – these being the only covariates collected across the CoRR study. While we acknowledge this limited covariate set is likely insufficient for formal harmonized analyses, it serves our exploratory purpose of illustrating the differences between causal and non-causal analyses. Moreover, even in mega-studies that collect more comprehensive covariate sets, many analyses focus primarily on age and sex for harmonization. We believe insufficient evidence exists to support the adequacy of such limited covariate sets for harmonization, and our work demonstrates how this limitation may lead to problematic confounding biases.

It is likely that other methods, such as extensions of our proposed `AIPW cComBat`, may be more appropriate in certain contexts than matching-based methods. In particular, for matching-based methods, one would typically restrict the study population (across all studies) to the “narrowest” covariate range; e.g., the “intersection” of the covariate distributions across the included studies. AIPW-based methods may instead present promise for aligning each dataset to a broader “reference”, and then sequentially estimating batch effects for each non-reference group against the broad reference group. This may be complementary to current efforts to produce so-called “lifespan” reference curves (Zhu et al., 2024), which propose harmonizing datasets on the basis of reference templates gradually developed over time. Our work highlights that the appropriateness of such comparisons require careful consideration of the demographic and phenotypic characteristics of both the groups used to develop these reference templates as well as the new datasets being aligned against the reference, which we believe has been insufficiently explored by the current literature. Further, AIPW-based methods generally offer the “doubly robust” property in parametric settings, where subsequent inference may be consistent if either the outcome or the propensity score model are correctly specified, affording two opportunities for correct estimation (J. M. Robins et al., 1994). Of note, our proof-of-concept `AIPW cComBat` implementation outper-forms `cComBat` and `cComBatGAM` approaches in simulation environments. However, these methods are more difficult to adapt to present approaches due to the fact that they necessitate the incorporation of weighting schemes to batch effect correction techniques. It is possible that there are other desirable characteristics (such as generalizability to broader target populations) in which AIPW-based methods are more optimal than matching-based methods, or alternative causal approaches that are more performant than those investigated here.

Our work highlights the question of how to properly interpret multi-site scientific studies, particularly regarding internal and external validity (Degtiar & Rose, 2023; Patino & Ferreira, 2018). When we re-weight data to impose demographic overlap, we can make internally valid conclusions in regions of the covariate space that support analysis, albeit potentially for a narrower set of covariate values. Our procedures learn a batch effect correction for a matched subset of individuals, with conclusions methodologically valid for individuals within a range of covariate overlap with this re-weighted population. This approach allows investigators to benefit from causal methods while minimizing discarded samples, balancing internal validity with statistical power for subsequent investigations. Less conservative approaches might attempt to apply the internally valid batch effect correction to a wider target population, requiring potentially dubious extrapolations of estimated batch effects to new covariate values. In such contexts, a more conservative approach would be to analyze data separately and derive conclusions across subsequent analyses (i.e., via meta-study).

While causal re-weighting procedures will attempt to impose demographic overlap (when possible), we explicitly avoid making recommendations as to sample sizes sufficient for subsequent inference. Rather, we believe that these balancing procedures should be considered in tandem with the proposed inference question (e.g., after correcting for batch effects), and subsequent conclusions described in terms of the retained sample. For instance, if one wishes to learn a relationship between age and brain connectivity across the lifespan from multiple datasets, but a “matched sample” only includes individuals between 20 and 25 across the different datasets, perhaps a meta-study would be more appropriate than a mega-study incorporating batch effect correction techniques. If one wished to continue with a mega-study approach, care should be taken that subsequent inference methods are sensitive to sample sizes (e.g., at a fixed effect size, a method would provide less confident answers with fewer samples) and that the inference is presented alongside descriptors of the retained sample (e.g., explicitly delineating that the analysis only applies to individuals between 20 and 25). Our software implementations, and consequently our simulations and real data analyses, default to raise errors if fewer than 30 samples are retained, a typical threshold for central limit theorem-based inference (such as regressions) (Rice, 2006), but this should not be used as a strict criterion.

Another limitation is that our methods exchange traditional statistical assumptions for sets of causal assumptions. This allows us to delineate a relatively understandable context (intuited via causal graphs and notions of covariate overlap) in which traditional batch effect correction techniques may be valid, and illustrate that data from mega-studies often do not resemble these assumptions. Even if “all models are wrong” (Box, 1976), their utility presupposes they loosely capture elements of the real data, which we show is frequently not the case for mega-studies.

### Future Work

This work suggests that harmonized analyses should be conducted with both harmonized measurement parameters and demographic distributions, departing from current practices in many megastudies (Di Martino et al., 2014, 2017; Yamashita et al., 2019; Zuo et al., 2014). Post-hoc batch effect detection and removal presents theoretical and conceptual inconsistencies regarding internal validity when not viewed through a causal lens. This is evident in the poor demographic overlap in popular neuroimaging studies like CoRR and ABIDE (Di Martino et al., 2014, 2017). While SRBPS (Yamashita et al., 2019) shows greater demographic overlap, both ABIDE and SRBPS introduce additional complexities by using neurodivergences in participant recruitment. If the measured connectome proxies underlying neurology, neurodivergent brain features may cause symptoms of different neurocognitive behavioral phenotypes (Hashem et al., 2020; Konrad & Eickhoff, 2010; Vogelstein, Bridgeford, Pedigo, et al., 2019), potentially introducing collider biases or differential-exposure measurement errors (Bareinboim & Pearl, 2012; Nebel et al., 2022; Pearl, 2014). This can be conceptualized as participant illness status yielding violations of the ignorability assumption (Bridgeford et al., 2023) and rendering some causal effects unidentifiable (Pearl, 1995, 2010a). While this problem has been defined for data fusion (Bareinboim & Pearl, 2016), its impact on batch effect detection or correction in high-dimensional biological studies remains unclear.

Recent work has proposed the use of deep learning methods for harmonization of magnetic resonance data derivatives such as T1w and T2w anatomical images (Hu et al., 2024; Liu & Yap, 2024). While to our knowledge no such developments have been proposed for fMRI, dMRI, or derivatives thereof (such as connectomes), we believe that these will likely be on the horizon. The methods proposed are heavily complementary to deep learning methods. In particular, deep learning methods are flexible for learning complicated batch effects from the data, and may hold similar promise to traveling subjects datasets for estimating and removing batch effects (Liu & Yap, 2024). However, deep learning methods are known be vulnerable to data dissimilar from that previously seen, known as the dataset or distribution shift problem, across many domains (Arjovsky, 2021; Ovadia et al., 2019; Taori et al., 2020). This is a particular concern for multi-site mega-studies, where as we have illustrated demographic overlap cannot be anticipated. We therefore do not anticipate that deep learning based batch effect correction methods will be immune to these concerns. We believe that incorporating many of these techniques with approaches such as vertex matching or other causal balancing procedures, as we propose in Section 2.2, may represent a future area of interest.

Future work will focus on applying these methods to correct batch effects in mega-studies like the Adolescent Brain Cognitive Development (ABCD) (Karcher & Barch, 2021), which includes *N* > 11,000 demographically diverse individuals across the United States, using a consistent, harmonized protocol. Studies may build upon the NKI-RS and Noble et al. (2017) by collecting measurements from diverse groups across multiple neuroimaging sites or protocols. Our work provides a theoretical foundation for evaluating studies like (Noble et al., 2017) and (Yamashita et al., 2019), which advocate aggregating multi-site *traveling subject* datasets to explore batch effects with minimal assumptions. This allows appreciation of internally valid demographic-specific effects and informs modeling assumptions, potentially enabling extrapolatory batch effect removal techniques in non-demographic-aligned datasets. Recent work highlights that data pre-processing strategies significantly impact subsequent derivatives (Bhagwat et al., 2021; Bridgeford et al., 2020; Gargouri et al., 2018; Vergara et al., 2017). While some strategies may optimally satisfy certain criteria (e.g., registration quality), they may improve or worsen batch effects. Future research could explore optimizing pre-processing to limit batch effects while meeting other requirements.

## Ethics Statement

This study analyzes publicly available neuroimaging data from the Consortium for Reliability and Reproducibility (CoRR) mega-study (Zuo et al., 2014). All data collection protocols from the original studies were approved by their respective institutional review boards (IRB), and data was shared through CoRR with approval or exemption from the original study IRBs and ethics committees. The research presented here involves secondary analysis of de-identified data (including de-facing of MRI data) and does not involve direct human subject experimentation. Our methodological approaches prioritize transparency and reproducibility through open-source code and detailed documentation. Care was taken to avoid over-interpretation of results, particularly when analyzing data with demographic imbalances or limited covariate overlap. When dealing with sensitive demographic variables like sex, age, and continent of measurement (as a proxy for race/ethnicity), we explicitly acknowledge the limitations of our covariate set and the potential for unmeasured confounding factors. The methods developed aim to promote more rigorous and ethical analysis of multi-site neuroimaging data by helping researchers identify when data may be insufficient to draw reliable conclusions, rather than potentially producing misleading results.

## Code Availability

The methods introduced in this work are made accessible for users via the `R` package `causalBatch` (Bridgeford et al., 2024) (available on `CRAN`), which features numerous package vignettes to facilitate ease-of-use. Code for reproducing the figures and analyses contained in this work is available from https://github.com/neurodata/causal batch, and a docker container with all software versions for reproducing these results is available at https://hub.docker.com/r/neurodata/causal batch.

## Data Availability

The raw data analyzed in this manuscript can be obtained at CoRR Mega-Study. The pre-processed data analyzed in this manuscript is available at http://neurodata.io/mri.

## Contributions

EWB wrote the manuscript; EWB and JTV revised the manuscript; EWB, JTV, and MP conducted study conception and design; EWB, JTV, BC, and MP conducted interpretation of results; EWB, JC, SP, and RL analyzed raw neuroimaging and demographic data; EWB and JTV devised the statistical methods; EWB performed statistical analyses; EWB wrote the software package; MP, GK, SN, TX, MM, BC, and JTV provided valuable feedback and discussion throughout the work.

## Acknowledgements

The authors are grateful for the support from the National Science Foundation (NSF) administered through NSF Career Award NSF 17-537, the National Institute of Health (NIH) through National Institute of Mental Health (NIMH) Research Project 1R01MH120482-01 and 1RF1MH123233-01, and the NIH through Research Project RO1AG066184-01. We also wanted to thank Betsy Ogburn for numerous insightful conversations regarding the presentation and framing of this work. We would also like to thank prior anonymous feedback for contributions to our manuscript.

## A Definitions

For a formal description of the below definitions, see Bridgeford et al. (2023). Informally, we adopt the following notation to describe batch effect correction. We assume that *Y*_*i*_ represents an observed measurement of interest (e.g., a connectome), *T*_*i*_ represents the batch in which the measurement is collected, *X*_*i*_ represents observed covariates, (e.g., age or biological sex), and *Z*_*i*_ represents unobserved covariates (e.g., height). The data (*Y*_*i*_, *T*_*i*_, *X*_*i*_, *Z*_*i*_) are *n* independent and identical samples from some true and unknown data generating distribution. The quantity 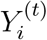 represents the measurement that would have been observed, had the measurement been collected in a given batch *t*. The key distinction is that *Y*_*i*_ represents the *actual* measurement, which is collected and studied. On the other hand, 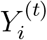 is a potentially hypothetical measurement, which is only actually observed for the case where *T*_*i*_ = *t* in most experimental contexts Cole and Frangakis, 2009. That is, individuals have a potential measurement 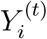 for every possible batch, but only a single of these potential measurements are studied under standard observational contexts. This is known as the *consistency assumption*, and can be written mathematically as:

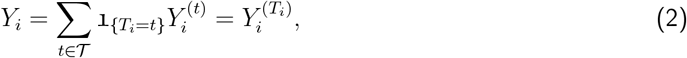

where *T* is the set of batch labels. That the observed measurements *Y*_*i*_ given the batch (or measured covariates) and the potential measurements 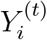 do not, in general, have similar distributions is known as the “fundamental problem of causal inference.” This problem requires resorting to additional assumptions to make conclusions on the basis of the observed data.

Using this convention, a batch effect is defined in Definition 1.

### Definition 1

(Batch Effect). *A batch effect exists between two batches t and t*^*′*^ *if* 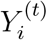 *and* 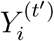 *havedifferent distributions*.

In light of this definition, a batch effect can be conceptualized as the *potential* measurements having different distributions between a given pair of batches. A batch effect is present if an individual’s potential measurements (which are random quantities) would differ (in terms of their distributions) *by virtue of them being measured* in batches *t* and *t*^*′*^.

The major conceptual leap is that, under the manner in which most mega-studies are collected (where individuals are *unique* to each batch), Definition 1 is about *potential* measurements rather than *realized* measurements. Causal claims regarding potential measurements will only be valid on the basis of the observed (realized) data insofar as the reasonableness of the assumptions upon which they rest.

### A.1 Associational Effects

In an associational context, we observe measurements *y*_*i*_ and batches *t*_*i*_ for each individual *i* ∈ [*n*], so effects can only be determined from realizations of *Y*_*i*_ and *T*_*i*_.

#### Definition 2

(Associational Effect). *An associational effect exists between batches t and t*^*′*^ *if Y*_*i*_|*T*_*i*_ = *t and Y*_*i*_|*T*_*i*_ = *t*^*′*^ *have different distributions*.

Likewise, we can use this intuition to develop a suitable definition for associational batch effect correction:

#### Definition 3

(Associational Effect Correction). *Associational effect correction is a function g, where:*

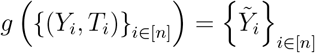

*such that for all pairs of batches t and t*^*′*^, 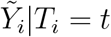 *and* 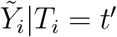 *have the same distribution*.

Whereas *Y*_*i*_ represents the measurement for an individual, the intuition is that 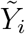 can be thought of as the “corrected” measurement, which does not have an associational effect for any pair of batches in the observed data.

Associational effects will often poorly characterize whether a batch effect is present. Consider, for instance, if in one batch *t*, we tend to measure older individuals, whereas in another batch *t*^*′*^, we tend to measure younger individuals. If age is related to the measurement we obtained, then the differences between *Y*_*i*_|*T*_*i*_ = *t* and *Y*_*i*_|*T*_*i*_ = *t*^*′*^ could be due to age or batch identity, and we have no way of differentiating whether the effect is a *bona fide* batch effect versus merely an associational effect. A sufficient condition for an associational effect to facilitate detecting or estimating a batch effect would be that individuals are randomized to each batch, in that individuals are randomly assigned to be measured in particular batches *pre-hoc*. Associational effect detection can be facilitated via Dcorr Székely et al., 2007, and associational effect correction can be facilitated via `ComBat` Johnson et al., 2007.

### A.2 Conditional Effects

In a conditional context, we observe measurements *y*_*i*_, batches *t*_*i*_, and covariates *x*_*i*_ for all individuals *i* ∈ [*n*], so effects can be determined from realizations of *Y*_*i*_, *T*_*i*_, and *X*_*i*_.

#### Definition 4

(Conditional Effect). *A conditional effect exists between batches t and t*^*′*^ *if for some covariate x, Y*_*i*_|*T*_*i*_ = *t, X*_*i*_ = *x and Y*_*i*_|*T*_*i*_ = *t*^*′*^, *X*_*i*_ = *x have different distributions*.

We can define conditional effect correction using this logic:

#### Definition 5

(Conditional effect correction). *Conditional effect correction is a function g, where:*

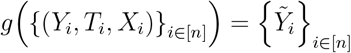

*such that for all pairs of batches t and t*^*′*^ *and for all covariate levels x*, 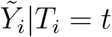, *X*_*i*_ = *x and* 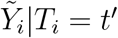, *X*_*i*_ = *x have the same distribution*.

Conceptually, given the consistency assumption from Equation 2, a conditional effect is equivalent to a batch effect if two conditions hold:

1. the measured covariates overlap in distribution between the batches (propensity overlap), and
2. the measured covariates provide all of the information regarding the mechanism about how people ended up in one batch versus the other (strong ignorability).

The former condition denotes that both batches must contain similar groups of people (in terms of measured covariates), and the latter condition specifies that the measured covariates *X*_*i*_ tell us all of the information needed to “exchange” measurements from one batch to the other. Borrowing the preceding example, even if we observe more young people in a batch *t*, we must still observe *some* young people in the other batch *t*^*′*^. In this sense, the measured covariates contain the information needed to “de-confound” disparities that might be batch effects or veridical effects due to upstream covariates. Therefore, when we make subsequent comparisons, we do not need to guess what people with similar covariates would have looked like in the other batch, and vice versa.

In this fashion, our comparisons can be thought of as locally (with respect to the covariates) exchanging a realized measurement *Y*_*i*_ in batch *t* with a realized measurement *Y*_*i*_ in batch *t*^*′*^, where both individuals have similar covariates *x*. Intuitively, these comparisons can therefore be conceptualized as synthetically comparing *Y* 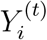 and 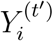 (the target estimand for establishing a batch effect) by using observed measurements *Y*_*i*_ from individuals who are similar on the basis of the covariates *X*_*i*_ between the two batches. Condition 1 ensures that we can make this intuitive step over the entire span of covariates in our batches. Conditional effect detection can be facilitated via cDcorr Wang et al., 2015, and conditional effect correction can be facilitated via `cComBat` Johnson et al., 2007.

In practice, we never know whether the propensity distributions overlap; we can only estimate them from the data. If our estimated propensities do not overlap given finite data, we again cannot reasonably differentiate between differences in the two groups being due to *bona fide* batch effects or empirical differences in the propensity distributions. This motivates a third approach.

### A.3 Adjusted Effects

As before, we observe measurements *y*_*i*_, batches *t*_*i*_, and covariates *x*_*i*_ for all individuals *i* ∈ [*n*], and we determine effects from realizations of *Y*_*i*_, *T*_*i*_, and *X*_*i*_. Prior to assessing the equality of any distributions, however, we weight the observations such that the observed covariate distributions are rendered approximately overlapping.

#### Definition 6

(Adjusted Effect). *An adjusted effect exists between batches t and t*^*′*^ *if after re-weighting samples such that X*_*i*_|*T*_*i*_ = *t and X*_*i*_|*T*_*i*_ = *t*^*′*^ *are approximately overlapping (or, alternatively, approximately equal) in distribution, Y*_*i*_|*T*_*i*_ = *t, X*_*i*_ = *x and Y*_*i*_|*T*_*i*_ = *t*^*′*^, *X*_*i*_ = *x have different distributions*.

We can similarly define adjusted effect correction as:

#### Definition 7

(Adjusted Effect Correction). *Assume that the samples are re-weighted via weights w*_*i*_ *(possibly* 0 *or* 1*) such that after re-weighting, X*_*i*_|*T*_*i*_ = *t and* 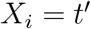 *are approximately overlapping (or alternatively, approximately equal) in distribution for all w*_*i*_*≠* 0.

*Adjusted effect correction is a function g, where:*

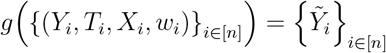

*such that for all pairs of batches t and t*^*′*^ *and for all covariate levels x*, 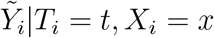 *and* 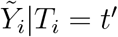, *X*_*i*_ = *x have the same distribution*.

Adjusted effects, by default, satisfy the first criterion for a conditional effect to be a batch effect. This is because rendering measured covariate distributions approximately equal intuitively is a more strict criterion than simply ensuring that they approximately overlap. The reason that we believe that reweighting to ensure approximate covariate distribution equality is desirable for effect correction, versus simply approximate covariate overlap, is discussed in Section 3.1 and Appendix D.3. Adjusted effect detection can be facilitated via Causal cDcorr Bridgeford et al., 2023, and adjusted effect correction can be facilitated with `Matching cComBat` (described in Section 2.3).

We still must satisfy the latter criterion for a conditional effect to be a batch effect; that is, given the measured covariates, we can ignore how people ended up in one batch versus the other. This assumption has the same interpretation as before.

### A.4 Crossover Effects

We observe measurements 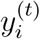 and covariates 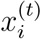, for all individuals *i* ∈ [*n*] and for all batches *t* ∈ *𝒯*. In this case, we can make statements on the basis of 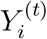 itself, because we actually observe outcomes for each possible batch. Therefore, we typically will not need to resort to local exchangeability (similar individuals/covariates across batches) as before, unless there are aspects of the individuals changing from one batch to another. *Crossover effects in general require the fewest assumptions to derive causal conclusions, since we directly observe all possible potential measurements for each individual*.

#### Definition 8

(Crossover Effect, constant states). *A crossover effect exists between batches t and t*^*′*^ *if, given that* 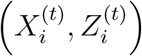 *and* 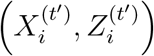 *are sufficiently similar*, 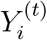 *and* 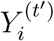 *have the same distribution*.

We are certain that any *traits* of the participants (i.e., variables that are constant for a given participant, such as genome) are the same across the two groups since the group members are identical (even if we did not measure these traits). However, *states* may differ as they may be functions of location or time. For example, if being measured impacts subsequent states, then a crossover effect may not be indicative of a batch effect without further assumptions and/or experimental design considerations (such as randomizing exposure order across participants, or resorting to adjusted effect strategies as before if these states are measured).

In the case where participant states are unchanging or are randomized, new associational strategies would need to be developed which, rather than comparing data directly across batches, batch effects would be estimated (or detected) by looking at disparities that arise across batches for the same individual measured multiple times. For instance, instead of investigating batch effects by comparing across batches, one could investigate batch effects by instead looking at within-individual differences across batches, and then investigating batch effects by aggregating across these within-individual differences.

In the case where participant states are changing and are not randomized but are measured, we can resort to adjusted strategies for adjusted effect detection or correction, via a crossover effect for non-constant states:

#### Definition 9

(Crossover Effect, non-constant states). *A crossover effect exists between batches t and* 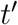 *if, after re-weighting samples such that* 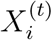 *and* 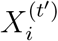 *are approximately overlapping (or, alternatively, approximately equal) in distribution, Y*_*i*_ ^*(t)*^ *and* 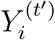 *have the same distribution*.

For this effect to be a true batch effect, we need to make the same assumptions as before; that is, that given the measured covariates (which include the changing states), we can ignore how people ended up in one batch versus the other. New methods would need to be devised which combine covariate adjustment strategies with similar strategies to those proposed to address crossover effects with constant states.

## B Statistical Methods

### B.1 Hypothesis Testing

Recall that statements of the form *f*(*y*) = *g*(*y*) against *f*(*y* ≠ *g*(*y*) are equivalent to *P*_*f*_ ≠ *P*_*g*_ against *P*_*f*_ *≠ P*_*g*_, as probability densities uniquely define distribution functions. Therefore, hypotheses for the effects described in Section 2.2 require two-sample and conditional two-sample testing procedures. A natural test statistic for the two-sample testing procedure is the Distance Correlation Székely et al., 2007, which is a non-parametric test for testing whether two variables are correlated. A simple augmentation of the distance correlation procedure Shen et al., 2017; Vogelstein, Bridgeford, Wang, et al., 2019 shows that `DCorr` can be used for the two-sample test, or a test of whether two samples are drawn from different distributions. `DCorr` is exactly equivalent in this context to the Maximum Mean Discrepency (MMD), which embeds points in a reproducing kernel Hilbert Space (RKHS) and looks for functions over the unit ball in the RKHS that maximize the difference of the means of the embedded points. When we instead consider the conditional two-sample test (i.e., a test of *f*(*y*|*x*) = *g*(*y*|*x*) against *f*(*y*|*x*) ≠ *g*(*y*|*x*)), we instead use the conditional distance correlation Wang et al., 2015, a kernel-based approach in which the points are embedded in a new, non-linear Hilbert Space, which augments the traditional linear Hilbert Space used in distance correlation to allow the definition of the squared distance covariance. Below, we let **Y** = (*y*_*i*_) ∈ *Y*_*n*_ denote realizations across both samples and 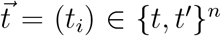 indicate from which sample each realization is drawn. **X** = (*x*_*i*_) ∈ *𝒳* _*n*_ denotes covariates which are known about the objects of interest.

#### Associational Effect

We have the following null and alternative hypotheses:

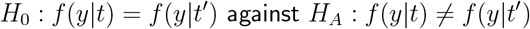

A test of the preceding hypotheses is performed using the distance correlation, and the natural test statistic is `DCorr` 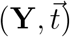.

#### Conditional Effect

We have the following null and alternative hypotheses:

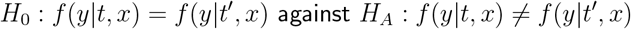

A test of the preceding hypotheses is performed using the conditional distance correlation, and the natural test statistic is `cDCorr` 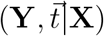.

#### Adjusted Conditional Effect

We have the following null and alternative hypotheses:

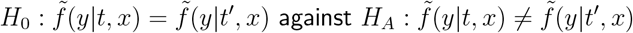

Unlike the preceding tests, we instead consider the data 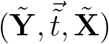, which are the measurements, sample indicators, and covariates of the *n* realizations after covariate adjustment. A test of the preceding hypotheses is performed using the conditional distance correlation, and the natural test statistic is `cDCorr` 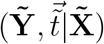.

#### Causal Crossover Effect

We have the following null and alternative hypotheses:

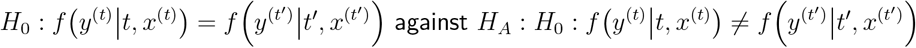

If the known covariates are identical between batches *t* and *t*^*′*^, we test the preceding hypotheses using the distance correlation, and the natural test statistic is `DCorr` 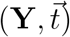. If the known covariates are not identical between batches *t* and *t*^*′*^, we test the preceding hypotheses using the conditional distance correlation, and the natural test statistic is `cDCorr` 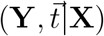. Note that to our knowledge, there are no energy statistical methods similar to other tests utilized herein that we are aware of for naturally paired data with multivariate responses that are non-parametric, so we use these tests as surrogates due to the lack of an alternative. These tests may afford robustness to certain types of model misspecification, at the expense of violations of independence assumptions across repeated samples. Therefore, the outcomes of such tests should be interpreted with caution.

#### More Than Two Batches

The above approaches generalize sufficiently to *K* batches using *K*-sample testing approaches Bridgeford et al., 2023. With *f*_*k*_ for *k* ∈ [*K*] denoting the densities associated with *K* sites or batches, this motivates hypotheses of the form:

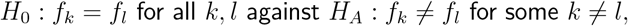

which can be tested using the distance correlation or the conditional distance correlation as above, with the caveat that 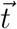 becomes the matrix **T** = (*t*_*i,k*_) ∈ {0, 1}_*n×K*_. Each entry *t*_*ik*_ = 1 if sample *i* is in batch *k*, and 0 otherwise. For the purposes of this manuscript, we focus on the two-batch case.

### B.2 Control Numerical Experiments

Control experiments are performed to ensure that after batch effect correction, the resulting data maintains interpretability and utility for scientific inquiry. Even if the data is devoid of batch effects, it must still be useful for downstream inference. For our connectome data, we investigate the preservation of demographic effects after batch effect correction.

#### Demographic Effect

Demographic effects are investigated across both the subset of connectomes upon which `Matching cComBat` is executed (the matched American Clique). We observe the tuple (*y*_*i*_(*k, l*), *s*_*i*_, *a*_*i*_, *t*_*i*_) for *i* ∈ [*n*], and *k, l* ∈ [*V*], where *V* = 116 denotes the number of parcels in the Automated Anatomical Labelling (AAL) parcellation N Tzourio-Mazoyer et al., 2002. We suppose that *Y* (*k, l*) is the [0, 1]-valued random variable denoting the weight of edge (*k, l*), *S* is the binary-valued random variable denoting the biological sex, *A* is the positive real-valued random variable denoting age, and *T* is the [*K*]-valued random variable denoting the batch. We let 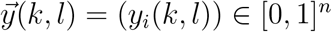 denote the realized edge weights,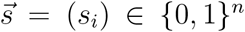 denote the realized biological sexes,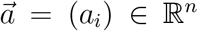 denote the realized biological ages, and 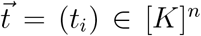 denote the realized batches. We say that a **demographic sex effect** exists when:

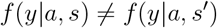

To test for a demographic sex effect, we have the following null and alternative hypotheses:

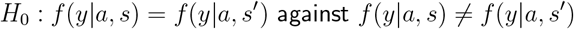

We are able to test the preceding hypothesis using the generalized covariance measure Shah and Peters, 2018, and the test statistic is 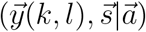.

### B.3 *p***-values and Multiple Hypothesis Correction**

*p*-values in this manuscript are estimated using permutation testing, which is an approach to obtain the distribution of the test statistic under the null with minimal assumptions and approximations Efron, 2004. All *p*-values are estimated using *N* = 1,000 (detection, Figure 6) or *N* = 10,000 (correction, Figure 7) permutations. Across all figures associated with this work, we are concerned with obtaining a proper estimate of the rate at which we detect effects (*discoveries*). Therefore, we control the false discovery rate (FDR) with the Benjamini-Hochberg Correction Benjamini and Hochberg, 1995.

## C Procedures for detecting and mitigating batch effects

### C.1 Detecting batch effects with Causal cDcorr

Many of the more direct types of detectable effects, such as associational and conditional effects, fail to adaequately account for confounding biases present in the data. We instead propose the use of Causal cDcorr, in which a conditional *K*-sample test (Wang et al., 2015) is performed on samples with the same “range” of covariate values after propensity trimming via a strategy known as vector matching (Lopez & Gutman, 2014). Specifically, from (Bridgeford et al., 2023), given a dataset with batch assignments *t*_*i*_, covariates *x*_*i*_, and measurements *y*_*i*_, Causal cDcorr is performed as follows:

1. Perform vertex matching, using the batch assignments given the covariate variables.
  a. Perform a multinomial regression, regressing the batch assignments *t*_*i*_ onto the covariates *x*_*i*_, to estimate a probability vector 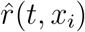 for each of the individuals for all batches *t* ∈ [*K*].
  b. For each batch *t*, use the procedure of (Lopez & Gutman, 2014) to produce high and low probability thresholds *l*^*(t)*^, *h*^*(t)*^.
  c. Exclude samples *j* for which 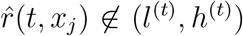 for any *t* ∈ [*K*]. This step excludes samples which are overly probable (or improbable) to be from any particular batch.
2. One-hot-encode the batch assignments *t*_*i*_ to obtain the *K*-dimensional vectors 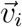, where *v*_*it*_ = 1 when *t* = *t*_*i*_ and 0 otherwise.
3. Compute the distance correlation between *y*_*i*_ and 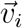 conditional on *x*_*i*_, using cDcorr (Wang et al., 2015).

This strategy is the focus of a complementary theoretical manuscript, in which we illustrate the theoretical and empirical (via simulations) benefits of this technique over competing approaches for detecting causal effects between potential outcomes. This strategy maintains both high sensitivity and specificity under traditionally problematic data regimes (high-dimensionality, non-monotonicities, and non-linearities) in which other methods typically fail (Bridgeford et al., 2023), making it a natural choice for causal discrepancy testing (of which “batch effect” detection, termed *causal unconditional discrepancy testing* in (Bridgeford et al., 2023), as-defined herein is a special case).

Figure 5C illustrates visually the causal procedures employed for adjusting the batches. Rather than fully matching to produce the adjusted batches, samples are retained only such that they have approximately overlapping covariate distributions (propensity trimming, shaded boxes). A full schematic illustrating the use of vector matching is detailed in Appendix E of (Bridgeford et al., 2023). In the event that the batch assignment mechanism is ignorable given the covariates and that effects between datasets are in the same direction across all covariate levels, the adjusted effect detected by Causal cDcorr is a causal batch effect, as proven in (Bridgeford et al., 2023). That the effects between datasets are in the same “direction” across all covariate levels can be best intuited via example. Consider the case where a batch effect exists, such that there is a signal disparity (a difference in the expected connectivity) in a particular edge of a connectome between two batches. If this signal disparity is a positive difference between batches 1 and 2 across all covariate levels (or, a negative difference across all covariate levels), Causal cDcorr will detect a causal batch effect. In the event that the assignment mechanism is ignorable but that effects between datasets are not in the same direction across all covariate levels, the adjusted effect detected by Causal cDcorr is a causal effect, but may reflect a *causal conditional discrepancy* (e.g., there are batch-specific differences, but they are isolated to particular covariate levels). In the event that the batch assignment mechanism is not ignorable, the effect may not reflect any causal effects (e.g., it may reflect unmeasured demographic differences between the batches).

### C.2 Mitigating batch effects using Matching cComBat

Unfortunately, many existing techniques for the removal of batch effects fail to adequately account for confounding biases that may be present in the data. We propose `Matching cComBat`, in which `cComBat` is performed on a subset of observational studies in which all pairs of studies are balanced against measured demographic covariates. `Matching cComBat` is performed as follows. Given measurements and covariates from *n* individuals across *K* batches, each of which has *n*_*k*_ individuals:

1. Perform vertex matching, using the batch assignments given the covariate variables.
2. Match control datasets to a reference dataset (defaults to smallest dataset).
  a. Perform nearest neighbor matching (without replacement) for all pairs of control datasets against the reference. This matching is performed many-to-one or one-to-many, with the aim of retaining the maximum number of possible matched pairs. When possible, use exact matching for categorical and binary covariates, and Mahalanobis distance matching for continuous and ordinal covariates. Default behavior uses a 0.1-width distance caliper (maximum possible distance for continuous and ordinal covariates).
  b. Retain all reference samples with a match to a control sample, and exclude all reference samples with no suitable matches (the matched reference samples).
  c. Retain control samples matched to a reference sample, and exclude control samples which are unmatched (the matched control samples).
3. Perform `cComBat` (Johnson et al., 2007) on the measurements of the matched reference and matched control individuals across the *K* batches, conditioned on the measured covariates *x*_*i*_.

In the event that the conditioning set closes backdoor paths (Pearl, 2009b, 2010b), `Matching cComBat` yields the removal of an internally valid causal effect and does not require extrapolation assumptions, unlike `cComBat` (Ho et al., 2011; Rosenbaum & Rubin, 1983, 1985; Stuart, 2010). If the conditioning set does not close backdoor paths, the effect removed is a conditional effect and may potentially yield the removal of demographic effects, as we saw in Figure 4. Appendix E.2 depicts the impact on the empirical covariate distribution of the adjustment procedure.

### C.3 Mitigating batch effects using AIPW cComBat

In addition to `Matching cComBat`, we propose `AIPW cComBat`, in which `cComBat` is combined with IPW methods to mitigate batch effects. `AIPW cComBat` is performed as follows. Given measurements and covariates from *n* individuals across *K* batches, each of which has *n*_*k*_ individuals, and a reference batch:

1. Perform vertex matching using the batch assignments given the covariate variables to remove samples with no overlap in covariate space.
2. Estimate propensity scores using multinomial logistic regression on the batch assignments given the covariates.
3. For each feature/dimension:
  a. Fit separate outcome regression models for each batch using the specified covariate model, and
  b. Calculate potential outcomes for each sample under each possible batch assignment using these models.
4. Compute the AIPW estimator for each batch and feature by:
  a. Weighting the difference between observed and modeled outcomes by inverse propensity scores, and
  b. Adding back the average modeled potential outcomes.
5. Adjust measurements by removing the estimated batch-specific component:
  a. Subtract the modeled outcome for the observed batch.
  b. Add back the modeled outcome for the reference batch.

This model provides double robustness, in that if either the outcome model for each batch or the propensity model are correctly specified, estimated batch effects are consistent for the true underlying batch effect, with respect to the reference batch.

### C.4 Covariate Adjustment

The exposed group *t* defaults to be the smaller of the two groups, and the unexposed group *t*^*′*^ is selected to be the larger of the two groups, where *n*_*t*_ is the number of individuals in the exposed group and 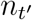 is the number of individuals in the unexposed group. The “covariate overlap” procedure attempts to ensure positivity of the propensity distribution for the unexposed group (i.e., *e*(*t*^*′*^|*x*) *>* 0). Intuitively, we exclude individuals from the unexposed group who do not appear “similar” to individuals in the exposed group. In this work, covariate overlap is established via vewrtex matching (Lopez & Gutman, 2017). Similarly, the “covariate balancing” procedure attempts to re-weight observations in the per-batch covariate distributions such that the covariate distributions are approximately equal (i.e., *f*(*x*|*t*) ≈ *f*(*x*|*t*^*′*^)). In this work, we perform *k* : 1 or 1 : *k* nearest-neighbor matching using the `MatchIt` package Ho et al., 2011. The number of matches 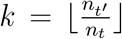 is chosen to be the largest number of unexposed matches possible. Individuals are balanced on the basis of individual sex, individual age, and continent of study. As there are likely many other categories with which brain connectivity may be confounded, we do not believe this covariate set is necessarily sufficient to identify a Causal Batch Effect 1, as we would need to be confident that these covariates exhibited the covariate sufficiency property. We perform exact matching on the basis of individual sex and individual continent of measurement, and use a 0.1-width distance caliper on the propensity score to obtain at most *k* matched participants for each treated individual Powell et al., 2020.

## D Simulations

### D.1 Batch Effect Detection Simulations

Simulations illustrating the sensitivity (high testing power when a causal effect is present) and specificity (tests which do not falsely detect effects) of `Causal cDcorr` for causal effect detection are in Bridgeford et al. (2023).

### D.2 Simple Simulations

This delineates the simulation settings for Figure 1 in our manuscript.

*n* = 500 points are sampled from Batch 0 or Batch 1 with probability 0.5; e.g., 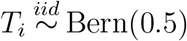.

#### D.2.1 Covariate distributions

In Figure 4, the covariate distributions are determined by:

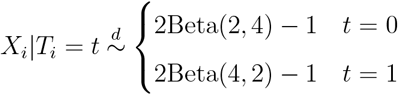

#### D.2.2 utcome model

With 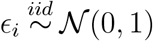, and the batch effect *β* is either 1 (bottom row) or −1 (top row):

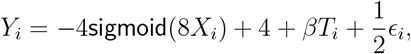

### D.3 Full Simulations

This delineates the simulation settings for Figure 4 in our manuscript.

*n* = 1000 points are sampled from Batch 0 or Batch 1 with probability 0.5; e.g. 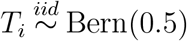.

#### D.3.1 Covariate distributions

In Figure 4, the covariate distributions are determined by:

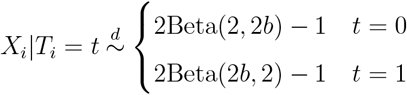

where *b* denotes the “unbalancedness”. We vary *b* from 1 to 5. Letting *f*_*0*_ denote the probability density function of *X*_*i*_|*T*_*i*_ = 0 and *f*_*1*_ the probability density function of *X*_*i*_|*T*_*i*_ = 1, the covariate overlap is:

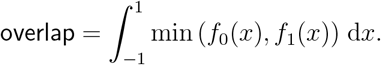

#### D.3.2 Simulation contexts

We investigate these in three simulation contexts, where for all simulations,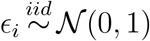, and the batch effect *β* is either −1 (Batch Effect) or 0 (No Batch Effect):

##### Non-linearity

A sigmoidal relationship between the covariate and the outcome. The outcome is:

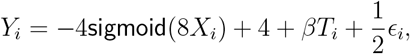

where sigmoid(*x*) is the non-linear sigmoid function; e.g.:

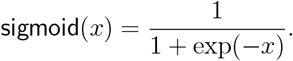

##### Non-monotonicity

A gaussian non-monotonic relationship between the covariate and the outcome. The outcome is:

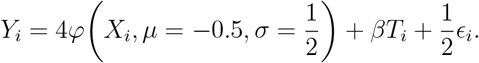

This non-monotonicity is “asymmetric” because *μ* = −0.5, which leads to the effect not being symmetric about *x* = 0 (whereas the covariates are, by construction, symmetric about *x* = 0).

##### Linear

A linear relationship between the covariate and the outcome. The outcome is:

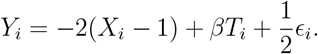

### D.4 Estimated Absolute Average Treatment Effect

To evaluate the effectiveness of each batch effect correction technique on simulated data, we compute the true data expected signal for each covariate level; e.g., 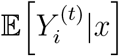 for each batch *t* and each covariate level *x*. Since *ϵ*_*i*_ has mean 0, this would be the quantity:

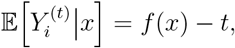

where *f* is the covariate/outcome relationship (possibly incorrectly modeled). Since *x* is continuous, we compute this value for *x* across *B* = 200 evenly spaced breakpoints for evaluation. Given a set of samples, we train a batch effect correction model, and fit the trained model to the expected signal for each covariate level, leaving us with corrected expected signal for each batch, which we denote by *v*(*t, x*). In theory, if the batch effect correction technique removes the batch effect as modeled, *v*(1, *x*) ≈ *v*(0, *x*) for all *x*. To evaluate each technique, we consider the estimated average absolute treatment effect (estimated AATE, for brevity) for each trial *r* of 1000 trials:

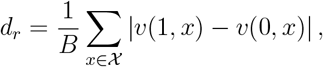

and the magnitudes of *d*_*r*_ are annotated in the plots (shaded red boxes) for a single simulation. We compute the mean estimated average absolute treatment effect (Mean Absolute ATE, Mean AATE) as:

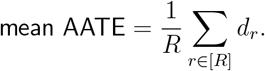

A value of 0 corresponds to the batch effect being completely eliminated (the expected signal for each batch after correction is identical), a value of 1 would equate to the AATE between the expected signals being the same as before batch effect correction, and a value *>* 1 corresponds to the residual disparity between the expected signals *exceeding* the original batch effect. The procedure for simulations is illustrated in Figure 8.

**Figure 8:**
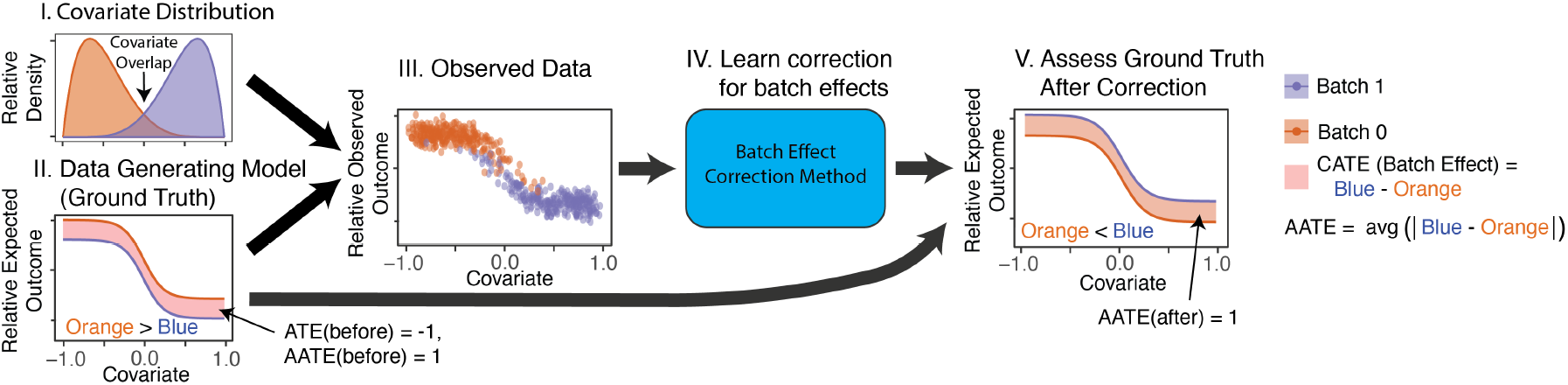
The procedure for assessing simulation performance in Figure 4. **I**. *n* individuals are sampled with equal probability from each batch. **II**. illustrates the generative model for the data in each batch. The ATE before correction is −1, and the average absolute treatment effect (AATE) is 1. **III**. illustrates the samples for each batch. **IV**. illustrates that the samples are used to train a batch effect correction model. **V**. the batch effect correction is applied to the data generating model, and the AATE is again computed between the two batches (and is still 1 here; e.g., the batch effect has not been removed). This procedure is repeated *R* = 1000 times for a given setting to produce the mean estimated AATE.

### D.5 No batch effect simulations

Figure 9A shows similar simulations to Figure 4, but now there is no batch effect whatsoever (the AATE before correction is 0). The goal of these simulations is to identify whether the different methods are able to identify the lack of a batch effect, and avoid introducing a batch effect. All the methods correctly estimate that there is no batch effect in the linear setting (Figure 4B.I). However, non-causal methods incorrectly estimate the presence of a batch effect for both the nonlinear and non-monotonic settings, except when the covariates are nearly perfectly overlapping (Figure 4B.II and Figure 9B.III), and introduce batch effects to the data. In these regimes, non-causal techniques tend to introduce a batch effect, when none is present *a priori*. The causal methods behave better here, correctly identifying the relative absence of a batch effect (and therefore avoiding the introduction of a batch effect).

**Figure 9:**
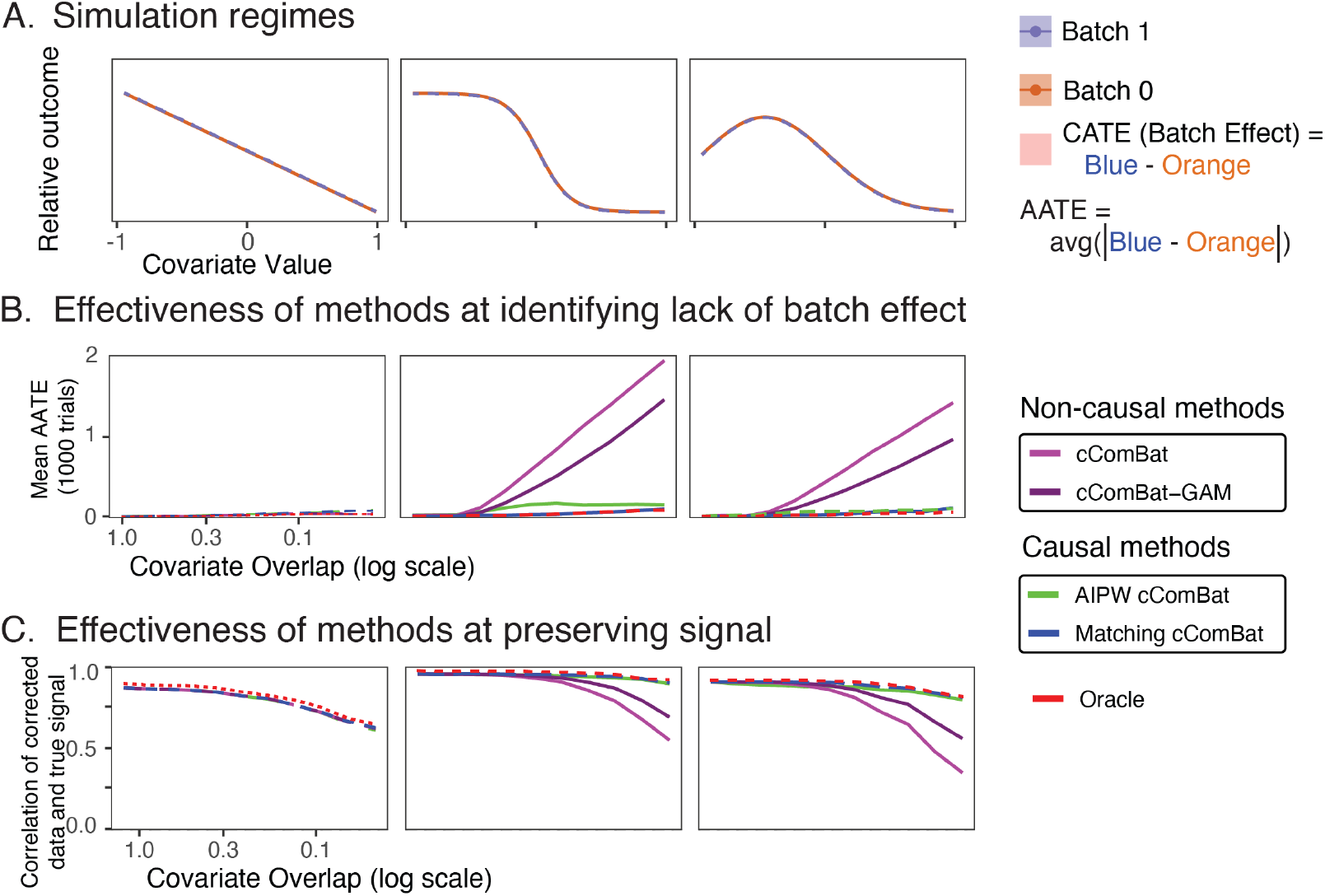
Simulation regimes illustrate that non-causal procedures are subject to strong biases without covariate matching. **(A)** illustrates the relationship between the relative expected outcome and the covariate value, for each batch (color), across **(I.)** linear, **(II.)** non-linear, and **(III.)** non-monotone regimes. The conditional average treatment effect (red box) highlights the batch effect for each covariate value. The average treatment effect (ATE) is the average width of this box, and the average absolute treatment effect (AATE) is the average absolute width of this box. In these simulations, the AATE before correction is 0. **(B)** The effectiveness of the techniques at avoiding the introduction of batch effects. Techniques with high performance will have a mean AATE after correction at or near 0 (no artifacts were introduced). **(C)** illustrates the effectiveness of different batch effect correction techniques for preserving the underlying true signal. Techniques with high performance will have higher correlations with the underlying true signal.

**Figure 10:**
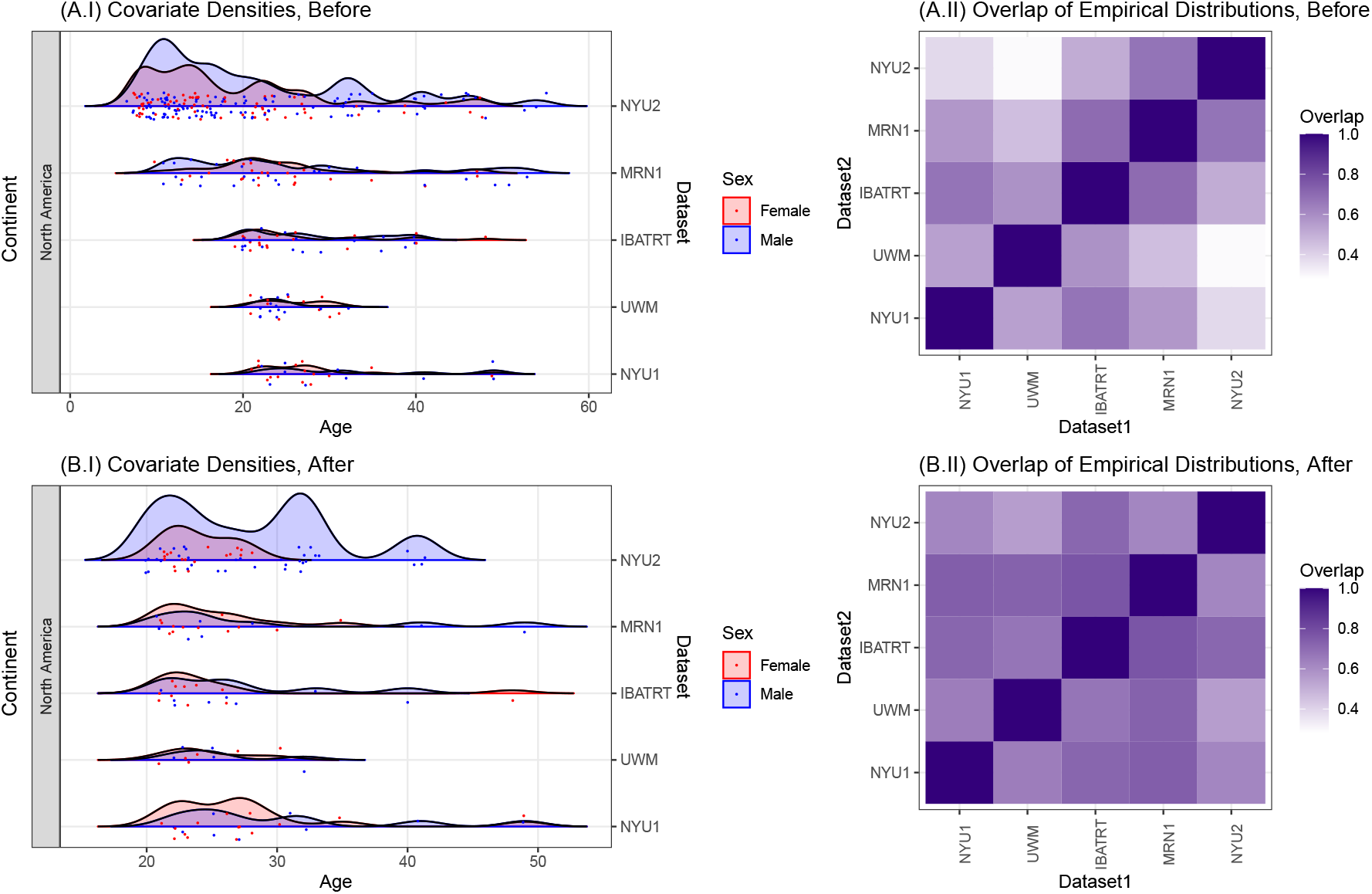
The overlap of the empirical covariate distributions for the American Clique, before and after adjustment. **(A)** The empirical distributions of the covariates before adjustment. **(B)** The empirical overlap of the covariate distributions after the adjustment procedure is applied, as discussed in Appendix E.2.

We evaluate how well the corrected data reflects the true underlying relationship (linear, non-linear, or non-monotonic) between the covariate and the outcome. We compare the corrected data to the true relationship using Pearson’s correlation Pearson, 1896, restricting to the matched samples so that the correlations are all computed with respect to the same set of sample in Figure 9(C). Low correlations indicate that the data poorly reflect the true relationship, suggesting that regardless of whether a batch effect were introduced to the data, the underlying signal has been perturbed. Causal methods here too tend to outperform non-causal methods for preserving the underlying signal in the data, and show performance near that of the `oracle`, particularly as covariate overlap declines.

## E Datasets

### E.1 Consortium for Reliability and Reproducibiliy (CoRR) data Pre-processing

The CoRR Mega-Study Zuo et al., 2014 is an aggregate dataset consisting of 27 studies collected with a similar goal: assessing the reliability and reproducibility of neuroimaging data. The mega-study consists of 1313 individuals, most of whom are measured numerous times, for a total of *N* = 3597 connectomes. All connectomes are estimated using the `m2g` (MRI to Graphs) pipeline Kiar et al., 2018, which provides a wrapper for the `CPAC` Pipeline Craddock et al., 2013. fMRI scans for each individual are first processed to remove motion artifacts using `mcflirt` Jenkinson et al., 2002. The fMRI scans are then registered to the corresponding individual’s anatomical scan using FSL’s boundary-based registration (BBR) via `epireg` Greve and Fischl, 2009. A non-linear transformation from the individual’s anatomical scan to the MNI152 Fonov et al., 2009 template is estimtaed using `FNIRT` Jenkinson et al., 2012. Nuisance artifacts are removed by fitting the voxelwise timeseries to a regression model incorporating regressors for the Friston 24-parameter model Friston et al., 1996, the top five principal components of the voxelwise timeseries in cerebrospinal fluid `aCompCor` Behzadi et al., 2007, and a quadratic drift term. The adjusted voxelwise timeseries is downsampled to the regions of interest (ROIs) of the Automated Anatomical Labelling (AAL) parcellation N Tzourio-Mazoyer et al., 2002 by taking the spatial mean signal for each timepoint across voxels within the region of interest. Functional connectivity is estimated using the pairwise correlation between all pairs of ROI timeseries within the AAL parcellation. Parcels are sorted throughout the manuscript according to hemispheric order, in which the parcels are aligned with left hemisphere parcels followed by right hemisphere parcels. Within hemisphere, parcels are sorted by AAL parcel number. For each study, we have baseline covariates for the continent, sex, and the age of participants.

#### The American Clique

The “American Clique” describes a subset of the CoRR Mega-Study in which the sample populations share similarities in sample demographic characteristics. These studies share a demographic focusing on males and females (in roughly equal proportions) of individuals across a wide age range, and include the “NYU2”, “IBATRT”, “MRN1”, “UWM”, and “NYU1” studies. The 833 connectomes comprising the studies of the American clique are reduced to the *N* = 284 connectomes with maximal demographic overlap identified through covariate adjustment C.4.

#### The NKI Rockland Sample

Nooner et al., 2012 is a single study from the CoRR Mega-Study consisting of 24 individuals, each of whom is measured two times across three functional MRI acquisition protocols, which vary in the repetition time for each slice of the sequence (TR). The data was collected with the intention of investigating the impact of the different MRI protocols in a crossover-randomized approach. Due to the crossover property, evidence in favor of an effect provides strong evidence of a causal batch effect. Images with a TR of 645 millisecond (ms), 1400 ms, and 2500 ms are measured, with the prompt for each subject remaining identical.

### E.2 Overlap of Empirical Covariates

The empirical overlap of the covariate distributions is difficult to compute in the case of data without making heavy parametric assumptions. For this reason, we turn to the distribution-free overlapping index Pastore and Calcagnì, 2019. The distribution free overlapping index,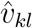 is computed by first approximating the density of the distribution of the measured covariates for each dataset *d, X* = (*A, S, C*), where *A* is a random variable whose realizations *a* ∈ *𝒜* are ages, *S* is a random variable whose realizations *s* ∈ *𝒮* are sexes (M or F), and *C* is a random variable whose realizations *c* ∈ *𝒞* denote continent, using the base `R` function `stats::density`. The random variable *D* has realizations *d* ∈ *𝒟* whose realizations denote dataset. The density *f*_*d*_(*a*|*S* = *s, C* = *c*) is the conditional density of age, conditional on the individual’s sex being *s*, continent of measurement being *c* for a given dataset *d*. The mass ℙ_*d*_(*S* = *s*|*C* = *c*) is the conditional mass of sex, conditional on the individual’s continent of measurement being *c*, for dataset *d*. Finally, the mass ℙ_*d*_(*C* = *c*) represents the mass of an individual’s continent of measurement being *c*, for dataset *d* (0 or 1 for all *d*, since all individuals from dataset *d* are either measured on continent *c* or not). An estimate of the overlap between the two densities, 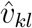 between datasets *k* and *l*, is computed using the formula:

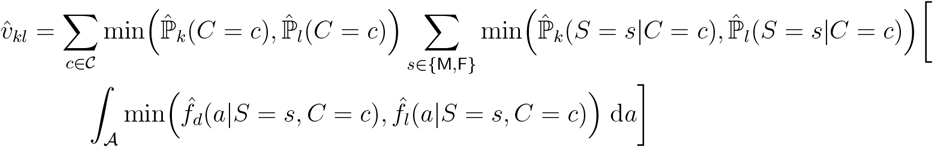

which is obtained via numerical quadrature.

Intuitively, this can be conceptualized as representing the mass of the “area under the curve” which is shared by the two densities for datasets *k* and *l*.

### E.3 Preservation of within-individual signal

The most fundamental properties of interest for bach effect correction methods to satisfy are that, for each individual, the connectomes after correction can be interpreted in the same context as the connectomes before correction. In this light, we investigate whether the topological properties of the connectomes are similar after correction as before. Figure 12(A) shows the connectomes before (Raw) and after batch effect correction is applied, by computing the cross-individual mean connectome. Note that before and after batch effect correction, the connectomes appear topologically similar, in that the relative edge-weights (across the methods) appear relatively consistent.

**Figure 11:**
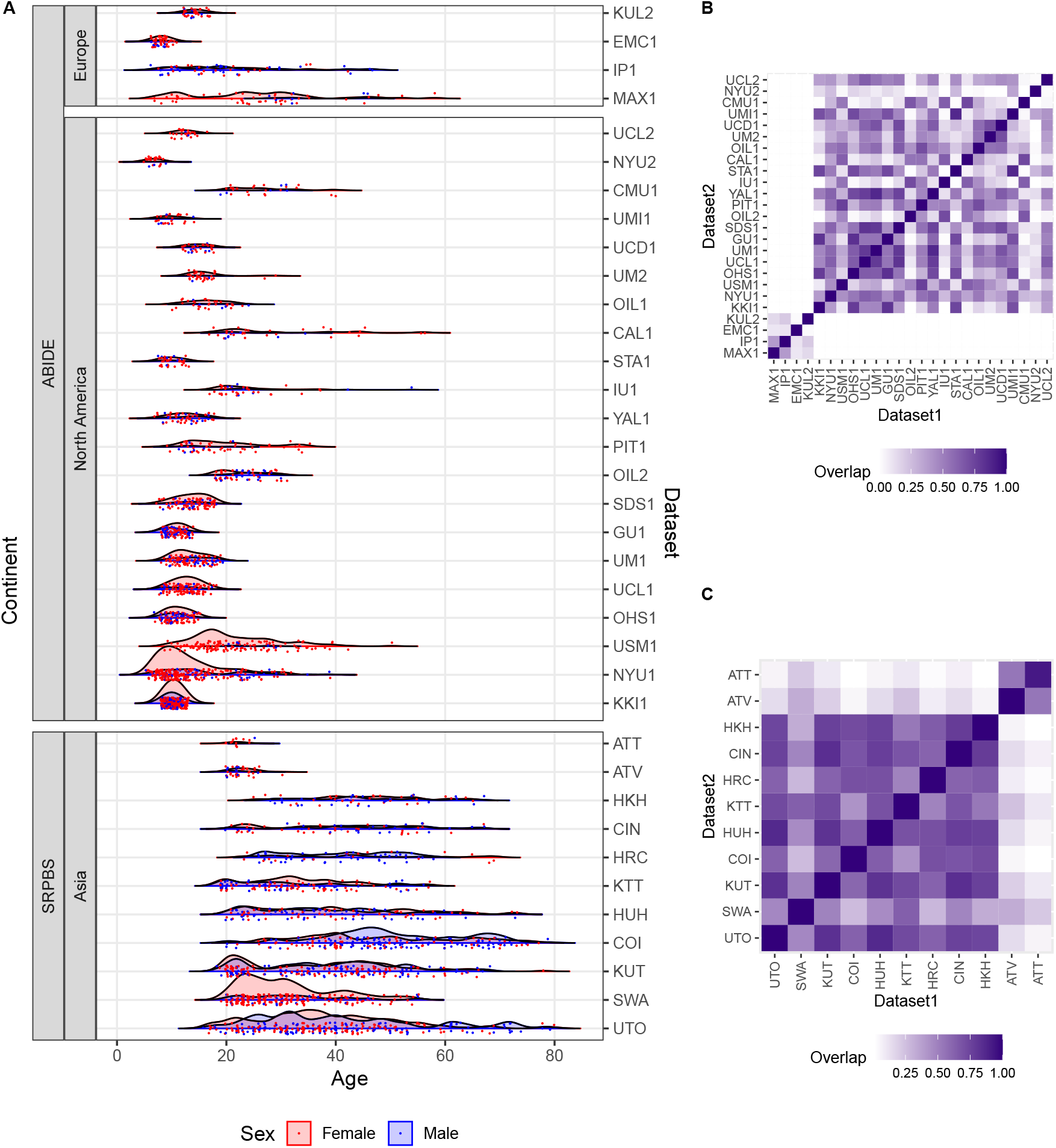
The overlap of the empirical covariate distributions for two mega-studies. **(A)** The empirical distribution of covariates. **(B)** The overlap of covariate distributions given by the distribution-free overlapping index for ABIDE mega-study Di Martino et al., 2014, 2017. **(C)** The overlap of covariate distributions given by the distribution-free overlapping index for the SRPBS mega-study. Yamashita et al., 2019. Like for the CoRR mega-study, while several pairs of sites have overlapping demographic distributions, many of the sites have extremely poor overlap in both mega-studies. In these cases, attempts to normalize for batch effects using model-based approaches like `cComBat` would be subject to the pitfalls of Figure 4 if modeling assumptions are not reasonable.

**Figure 12:**
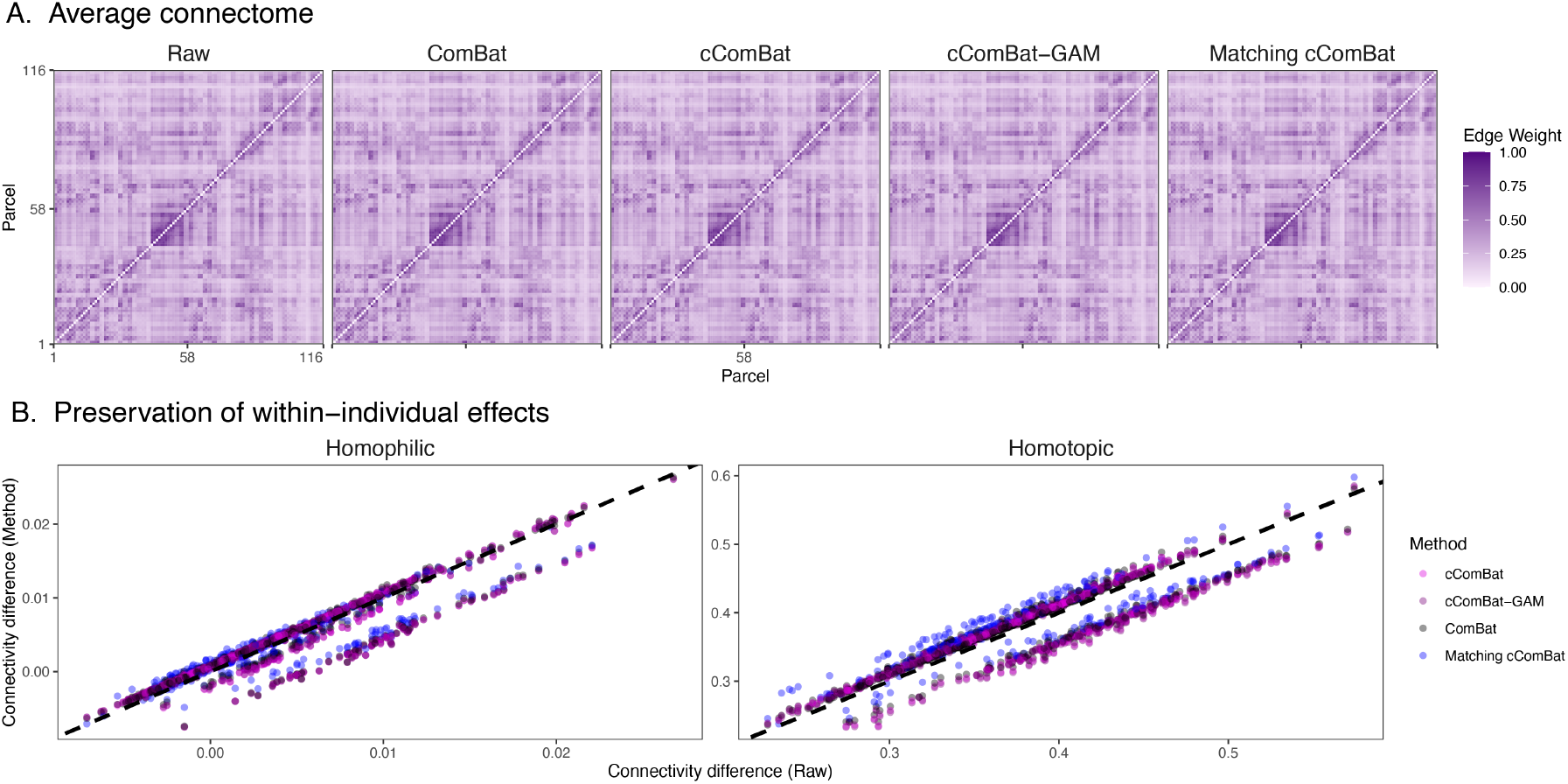
Preservation of Topological Properties of Connectomes after Batch Correction. **(A)** The average connectome, before (Raw) and after batch effect correction (other columns) across all individuals in the American Clique. **(B)** Scatter plots of two topological features of connectomes, homophily and homotopy. The *x-axis* denotes the average edge weight between edges which satisfy the noted feature and edges which do not, before any correction is applied (the “raw” connectomes). The *y-axis* denotes the same property for connectomes after correction is applied (point color).

Figure 12(B) considers two properties of functional connectomes, homotopy and homophily. Homotopic edges are edges between ROIs in the same hemisphere of the brain (e.g., two an edge between two nodes in the left hemisphere). Homophilic edges are edges between ROIs which denote the same brain area, but are in opposite hemispheres of the brain (e.g., an edge between the left and right motor cortex). In general, functional connectomes show a slight homophilic effect, and a very strong homotopic effect (Chung et al., 2020). We compute the effect size for each individual before and after correction, as the difference in average connectivity for edges in the noted edge group and edges not in the noted edge group (e.g., a comparison between the average connectivity of homophilic and non-homophilic edges, for “Homophilic”, and a comparison between the average connectivity of homotopic and non-homotopic edges, for “Homotopic”). Points falling along the diagonal dotted black line *y* = *x* tend to have a similar signal effect before and after batch correction, which includes the vast majority of the individuals.

